# VAMP5 is a novel target for selective inhibition of gliomas with high NDRG4 expression via downregulating PLK1 and beyond

**DOI:** 10.1101/2025.05.12.653623

**Authors:** Yinghao Zhang, Haoran Wang, Jinhai Wang, Yufang Sui, Wanhong Zhang, Yisheng Liu

## Abstract

**Background:** SNAREs participate in cancer’s progression or stress resistance. But the current conclusions on their involvement in cancers are generally unsystematic and even contradictory on one SNARE’s roles in different types of cancer. It’s necessary to find universal mechanisms of SNARE’s participation in cancers possessing generalizability within appropriate range for application of targeted therapy.

**Methods:** Our study initiated with database analysis on clinical information and bulk RNA-seq data to locate potential effective target for certain cancer type. Then experiment validations/explorations were performed on clinical human sections, cell lines and xenograft models. Lentivirus (LV) were applied for gene KD, overexpression and microRNAs sponge; Bulk RNA-seq, microRNA-seq and DIA proteomics for identification, quantification, genesets/target analysis; WB, single-IF, TSA (Tyramide signal amplification) multi-IF, IHC for proteins detection using antibodies; QPCR for artificial microRNA-sponge sequences test; Other reagents/instruments applied for corresponding purposes.

**Results:** VAMP5 is a therapeutic target for gliomas with expression selectivity. Gliomas could be divided into 2 types according to effects after VAMP5 KD. In VAMP5-KD sensitive type, PLK1, a classical gene vital for tumor growth/survival, is universally protein-level down-regulated that mediated partly via decrease of several up-reg microRNAs like mir-1301-3p and mir-12135, causing growth inhibition. While among insensitive type that VAMP5-KD is invalid for growth inhibition, PLK1 is not altered consistently, which serves as an evaluation marker for inhibition effectiveness. NDRG4 is a prediction marker for VAMP5-KD sensitivity that sensitive gliomas express higher NDRG4 than insensitive type on RNA level. Moreover, NDRG4 overexpression converts VAMP5-KD insensitive gliomas into sensitive type and additionally cause protein-level downregulation of VCAM1 and especially TNKS which also benefit the growth inhibition and PLK1 reduction partly. Except for results above, LGALS1 correlated and colocalized with VAMP5 possessing broader consistency than other membrane proteins so indicates the immune-enhancing potential of down-reg LGALS1 via KD VAMP5.

**Conclusions:** These discoveries elucidated the mechanism of VAMP5 KD on gliomas inhibition preliminarily. New sights for glioma therapies including selective therapeutic target, prediction marker, evaluation marker, enhancing strategy and correlated immuno-checkpoint markers were concluded with further development potentials.

## Introduction

SNARES (Soluble *N*-ethylmaleimide-sensitive factor attachment protein receptors) are a class of proteins related to membrane-fusion based secretion, endocytosis and intracellular transportations that involved into many biological processes. The most classical example is vesicle-membrane fusion process of neurotransmitter release from presynaptic terminal. Others like TCR-activated cytotoxic T lymphocyte secretion of lytic granules [1–3], secretion of atopic dermatitis related cytokines in human keratinocytes [4] and the membrane translocation of ion channels TRPV1, TRPA1 in sensory neurons [5]. As an abnormal state, there must be biological processes related to membrane fusion disorders in tumor cells, therefore SNAREs may be linked to oncogenic process, which have been supported with many clues. YKT6 promotes the migration and invasion of oral squamous cell carcinoma and non-small cell lung cancer cells, respectively [6, 7]; STX12 and SNAP23 mediate membrane trafficking of SRC, EGFR and ITGB1, facilitating the formation of invadopodia in breast cancer and fibrosarcoma to support migration/invasion [8]. Similarly, SNAP23, VAMP7, STX4 mediated the membrane trafficking towards invadopodia and secretion of matrix-metalloproteinase supporting breast cancer invasion [9].VAMP8 coupled with Rab17 confine the pro-invasive factor within endosome and lysosome, meanwhile knocking down of VAMP8 strengthen pro-invasive factors secretion and invasiveness of breast ductal carcinoma [10]. STX6 mediates the endosomal transportation of mutated EGFR from plasma membrane to nucleus translocation and promote proliferation of glioblastoma multiform [11, 12]. STX6 together with SMPD1 are indispensable for plasma membrane translocation of MET, a membrane protein of RTKs that are often overexpressed and oncogenic in several cancers [13]. SNAP23, SNAP29 and VAMP3 jointly mediate the movement of KRAS from endosome to correct location of plasma membrane followingly transducing signals supporting tumor development [14]. STX3 mediates the ubiquitination of PTEN related to tumor suppression and its subsequent degradation to promote tumor growth via PI3K-AKT-mTOR signaling in breast cancer [15]. The autophagosome-lysosome fusion process mediated by VAMP8, SNAP29 and STX17 [16] is associated with several stress resistance of cancers like TRPM7-agonist-stimulating induced tumor suppression [17], a new drug S670 induced ferroptosis in glioblastoma [18] and Temozolomide treatment in same type of cancer [19].

Belonging to SNAREs, VAMP5 was firstly identified at 1998 preferentially expressed in heart and skeletal muscle but not in brain [20]. However, the subsequent researches on this gene were obscure and non-systematic. At 2010, VAMP5 was demonstrated not mediate the membrane fusion with t-SNARE complexes including syntaxin1/SNAP-25 and syntaxin4/SNAP-25 in vitro [21]. At 2013, its localization were proved again with experiments mainly at cardiac and skeletal muscle and not in brain and intestine [22]. In 2016, sequence variations of VAMP5 was confirmed related to the risk of total colonic aganglionosis [23]. In 2018, the vital roles of VAMP5 in urinary and repository systems were confirmed via gene knocking-out mice, in which depletion of VAMP5 showed higher rate of embryo death accompanied by insufficient expansion of lung, and the viable mice showed ureter duplication [24]. In 2020, a pure bioinformatic result proved 10 genes including VAMP5 were most suitable input factors for random forest model predicting tumor purity of low-grade-glioma RNA-seq samples [25, 26]. In 2022, VAMP5 was demonstrated secreted in extracellular vesicles of mice retinal Muller glia cells responsive to ischemia [27]. In recent 2024, 2 researches were relevant to VAMP5 applying bioinformatic analysis plus experiment validations. The former found 5 marker genes including VAMP5 that were high expressed in tumor-specific T cells related to immune response to bladder cancer positively [28]; The latter identified 5 marker genes including VAMP5 in T cells representing kidney transplantation rejection [29]. Except for these clues, a review in 2024 also mentioned selective expression of VAMP5 in low grade glioma but without further discussions [30] . Straightly to say, no experimental conclusions of VAMP5 on its direct participation in cancers.

SNAREs’ involvement in cancers and reviews on VAMP5-related researches were generally incoherent and unsystematic. Some present results mentioned above on SNAREs in cancers were even contradictory like the tumor supporting/inhibiting roles of VAMP8 and the opposite transportation directions of STX6 and EGFR in different cancers. It seems that roles of one SNARE gene can be different in various types of cancers. Our research initiated via prior-knowledge independent bioinformatic analysis on large database samples to explore effect SNARE therapeutic targets possessing universal commonality within broader range, and followingly validated with experiments to reveal the mechanism.

## Results & analysis

### VAMP5 is a promising target for gliomas therapy with tumor expression selectivity

The GSEA workflow on TCGA samples was overviewed in fig 1A. All the cancer type participated in GSEA analysis listed in table 1. Only “brain glioma” and “adrenal gland adenoma & adenocarcinoma” type showed significant enrichment of SNARE gene sets (elements shown in table S1). VAMP5, with simple splicing structure (fig 1e), showed largest foldchange in GSEA core enrichment genes, satisfying survival improvement associated with its downregulation, preferential expression in gliomas clinical samples or cell lines compared to normal brain cells/tissues and especially between GBM and paracancerous tissues within same patient (fig 1b, 1d, 1f-h). Meanwhile VAMP5 highly correlated with several genes that positively influence tumor growth or immune escape among hundreds of TCGA samples (fig 1c) like TMSB4X [31], CLIC1 [32–34], LGALS1 [35–41], VAMP8 [16, 19], RRAS [42], ARPC3 [43] and TMSB10 (predicted by sequence similarity). Therefore, descending VAMP5 may in turn down-regulate these genes and reciprocally possible for tumor suppression with more universal range. Another cancer type that SNAREs were significantly enriched, Adrenal gland cancers (GSEA result shown in fig S1a) are overall easy for diagnosis and surgical removal so not focused for subsequent investigation. Simultaneously, compared to VAMP5, other core enrichment genes in gliomas type, like SEC22B that with least P value between withtumor/tumorfree team (fig 1b) did not behave better on survival improvement or selective expression (fig S1b-1c) so were not concerned in this research either. Though not chosen, these potential directions also merit profound investigations.

**Fig 1:**
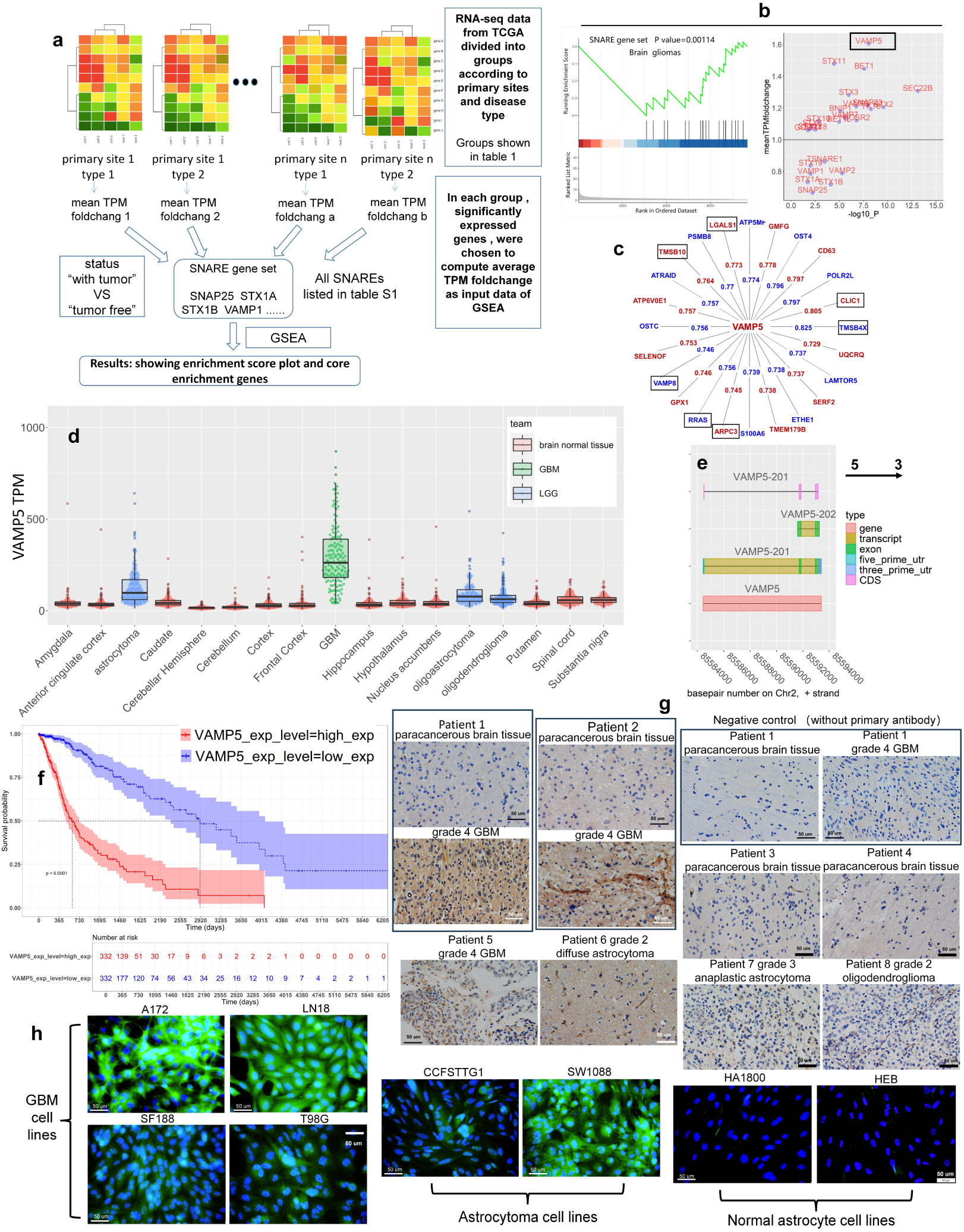
VAMP5 is a potential target for glioma treatment according to database analysis and experiment validation. a: GSEA working flow on TCGA samples: b: GSEA enrichment score plot of brain glioma samples in TCGA and volcano plot of core enrichment genes; c: Genes correlated with VAMP5 according to WGCNA among TCGA samples in brain glioma type; several genes published supporting caners’ survival or development were marked in black rectangles; d: Box-beeswarm plot of VAMP5 expression in TCGA coupled with GTEX samples; e: Gene track plot of human VAMP5 visualized according to ensemble gene annotations; f: Survival analysis result of VAMP5 among TCGA samples; g: IHC staining VAMP5 on clinical paraffin sections; h: Immune fluorescence results staining VAMP5 on 6 glioma cell lines, A172, T98G, LN18, SF188, SW1088, CCFSTTG1 and 2 immortal astrocyte cell lines HA1800 and HEB.

**Table 1:**
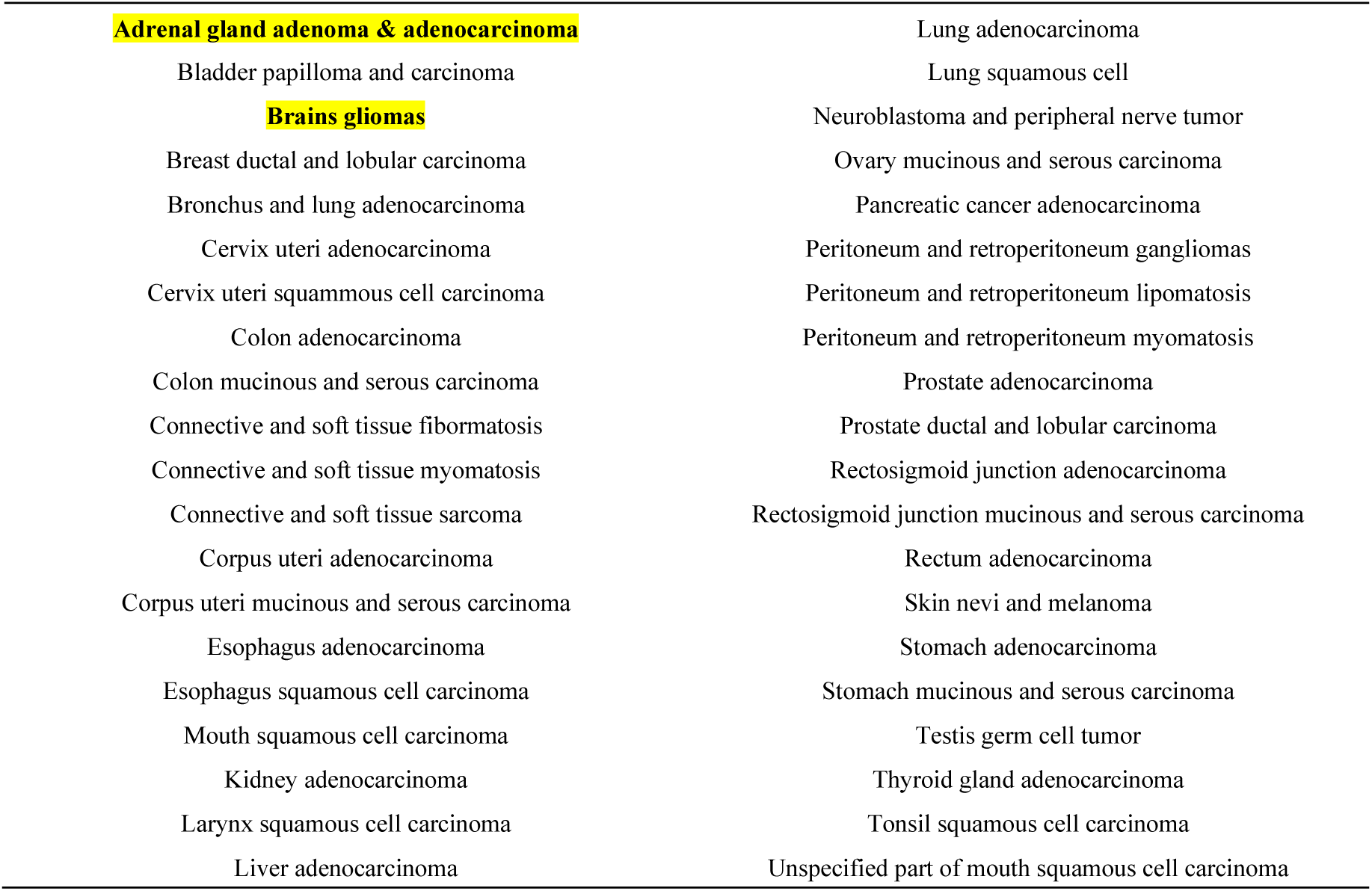
TCGA transcriptome grouped by primary site and disease type; RNA-seq data of all types were taken into GSEA computation as figure 1A indicated; Those types that SNARE gene set was significantly enriched were marked in yellow.

|  |  |
| --- | --- |
| **Adrenal gland adenoma & adenocarcinoma** | Lung adenocarcinoma |
| Bladder papilloma and carcinoma | Lung squamous cell |
| **Brains gliomas** | Neuroblastoma and peripheral nerve tumor |
| Breast ductal and lobular carcinoma | Ovary mucinous and serous carcinoma |
| Bronchus and lung adenocarcinoma | Pancreatic cancer adenocarcinoma |
| Cervix uteri adenocarcinoma | Peritoneum and retroperitoneum gangliomas |
| Cervix uteri squamous cell carcinoma | Peritoneum and retroperitoneum lipomatosis |
| Colon adenocarcinoma | Peritoneum and retroperitoneum myomatosis |
| Colon mucinous and serous carcinoma | Prostate adenocarcinoma |
| Connective and soft tissue fibromatosis | Prostate ductal and lobular carcinoma |
| Connective and soft tissue myomatosis | Rectosigmoid junction adenocarcinoma |
| Connective and soft tissue sarcoma | Rectosigmoid junction mucinous and serous carcinoma |
| Corpus uteri adenocarcinoma | Rectum adenocarcinoma |
| Corpus uteri mucinous and serous carcinoma | Skin nevi and melanoma |
| Esophagus adenocarcinoma | Stomach adenocarcinoma |
| Esophagus squamous cell carcinoma | Stomach mucinous and serous carcinoma |
| Mouth squamous cell carcinoma | Testis germ cell tumor |
| Kidney adenocarcinoma | Thyroid gland adenocarcinoma |
| Larynx squamous cell carcinoma | Tonsil squamous cell carcinoma |
| Liver adenocarcinoma | Unspecified part of mouth squamous cell carcinoma |

### Glioma cells could be divided into VAMP5-KD sensitive and insensitive type that VAMP5-KD resistance should be noticed

Real VAMP5 KD experiments with lentivirus (LV) were performed on several glioma cell lines. 4 cell lines, CCFSTTG1, SW1088, A172, SF188 were validated sensitive to VAMP5 KD that their growth were significantly inhibited (fig 2a-d, 2f, fig S2) and defined as “VAMP5-KD sensitive type”. Simultaneously, another 4 glioma cell lines, LN18, T98G, HS683 and U87 were defined as “VAMP5-KD insensitive type” that adequately KD VAMP5 will not inhibit growth (fig S3). 2 cell lines belonging to sensitive type, CCFSTTG1 and SF188 showed resistance to VAMP5 KD and require more KD lentivirus addition to overcome this hinderance, while in other sensitive cell line like SW1088 and A172, MOI 10 is enough for VAMP5 KD (fig 2e and fig S2). We did not observed distinctions on tumor migration or invasion after VAMP5 KD, meanwhile secretion of VAMP5 was not tested via WB on conditioned medium either (data not shown). Therefore, these characteristics were not concerned.

**Fig 2:**
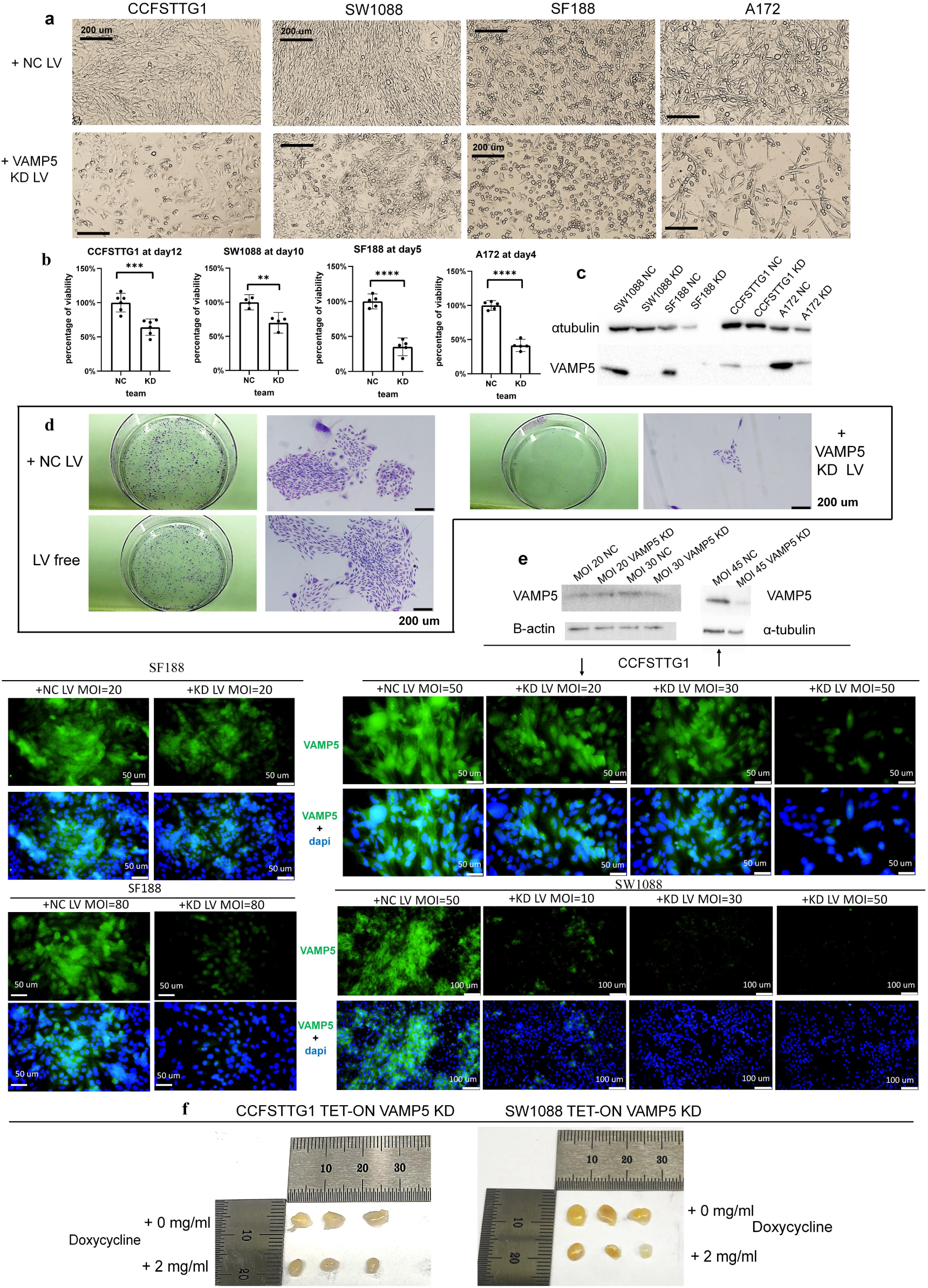
VAMP5 KD was validated effective for part of glioma cell lines growth inhibition; a: Bright field living images of 4 VAMP5-KD sensitive cell lines; b: Cell viabilities of 4 sensitive cell lines; c: WB result; d: Colony formation assay of one cell lines CCFSTTG1; e: WB and IF results indicating the resistance effect of VAMP5 KD in cell lines CCFSTTG1 and SF188, compared to VAMP5-KD unresistant SW1088; f: in vivo tumor sizes comparation of SW1088 and CCFSTTG1.

### VAMP5 KD among sensitive type induced universal down-regulation of G2/M cell cycle-related genes and caused virus-response related gene sets up-regulation in KD resistance cell lines

We believe a system including same cancer type, input VAMP5-KD condition and outcome of growth inhibition tends to work through same pathways, thus RNA-seq and followingly GSEA searching enriched gene sets intersections of MsigDb was performed, results shown in fig 3. Jointly enriched gene sets were not found in C5 gene sets (data not shown). However, “Fischer G2/M cell cycle” [44] (FGCC) was enriched with maximum consistency in C2 sets that only failed in SF188 (fig 3a). Meanwhile, several gene sets linked to upregulation response to virus infection were also discovered in VAMP5-KD resistant SF188 and CCFSTTG1 but not in SW1088 and A172 that did not show apparent resistance. It should be noticed that though cells were clearly inhibited, classical death pathways like necrosis, apoptosis, pyroptosis, autophagy or ferroptosis were not RNA-level enriched though these gene sets recorded in MsigDb and further apoptosis verification testing caspase3/7 activity didn’t find apparently positive result either (fig S4). Heatmaps of FGCC related genes were visualized (fig 3b), in which KIF20A was observed downregulated universally among cell lines (fig 3c). Moreover, a published research on medulloblastoma, a cancer type similar to gliomas, also demonstrated the apoptosis-independent inhibition when KIF20A downregulated [45], which was consistent with our result. Additionally, PLK1 is an upstream interactor and phosphor-regulator of KIF20A [46] though it behaved incomplete consistency of down-regulation compared to KIF20A. Consequently, these 2 proteins were candidating targets for next step.

**Fig 3:**
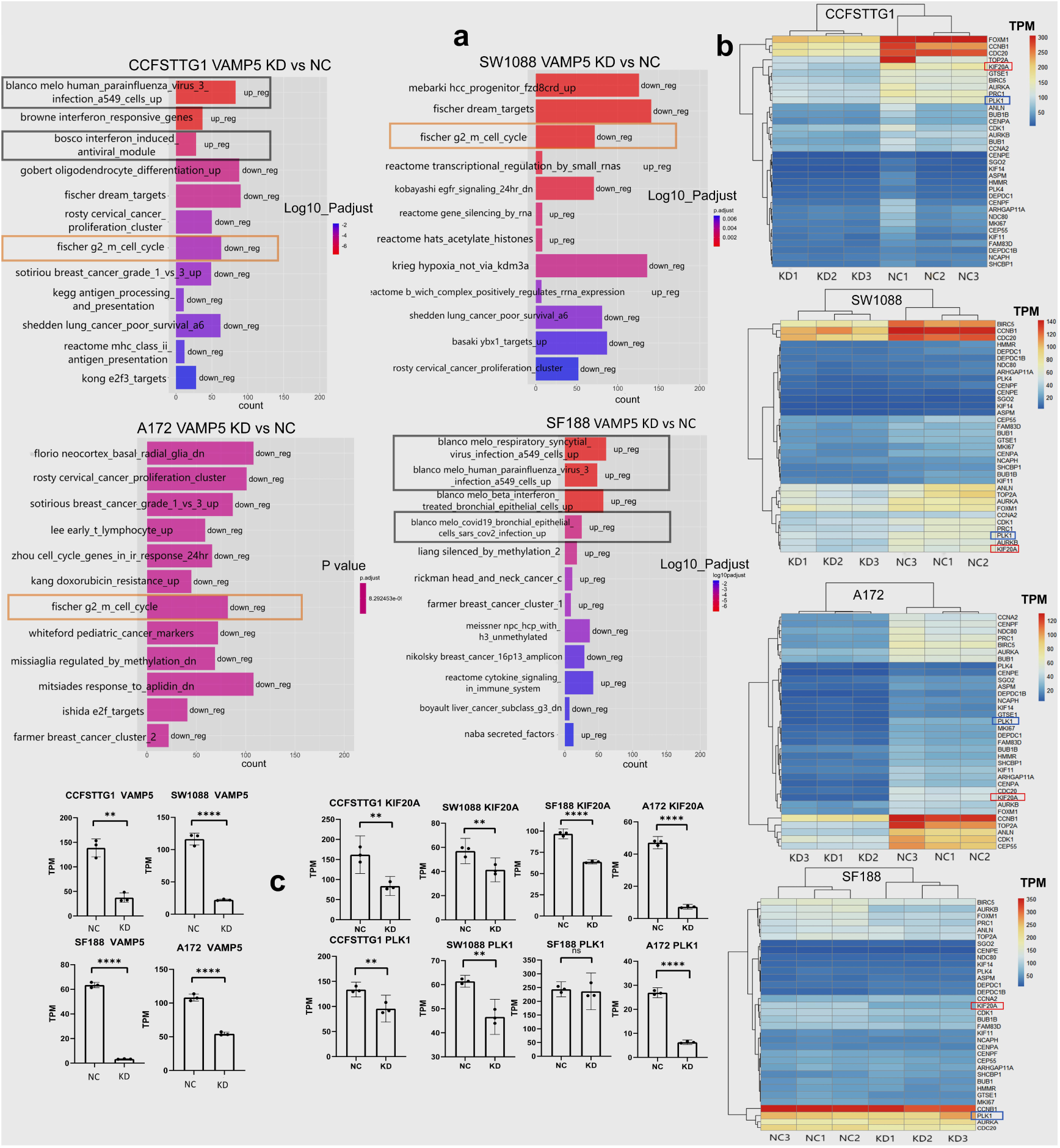
RNA-seq analysis result of 4 VAMP5-KD sensitive cell lines; a: GSEA result searching MsigDb C2 sets, in which 12 representative enriched gene sets were exhibited. The overall consistent enrichment sets were marked in orange rectangles, and those sets that may related to VAMP5-KD resistance were marked in black rectangles; b: The cluster heatmaps indicating union set expression of core enrichment genes belonging to “Fischer G2/M cell cycle” set among 4 cell lines; c: TPM barplot expression of KIF20A, PLK1 and VAMP5.

### PLK1 is a better evaluation marker to distinguish VAMP5-KD sensitive/insensitive type of gliomas than KIF20A and demonstrated responsible for induced growth inhibition

Immune fluorescence (IF) performed on KIF20A and PLK1 combining semi-quantification (fig 4a-d). KIF20A downregulated after VAMP5 KD in 2 of 4 sensitive cell lines but not lowered apparently enough in A172 and not changed in SF188, while PLK1, a classical cancers treatment target and G2/M cell cycle regulator [47–50], showed perfectly consistent lower-down among 4 cell lines. PLK1 behaved better on alteration consistency than KIF20A. The G1/G2 transition disorder phenomenon was proved via cell cycle test (fig 4e), in which G1 peaks were lowered while death cell rather than G2/M peaks elevated. As comparative, PLK1 expression was not altered in all 4 insensitive glioma cell lines (fig S5). Growth facilitating functions of PLK1 was validated via its inhibitor and degradation inducer, GSK461364A among sensitive cell lines CCFSTTG1, SW1088, insensitive type T98G, U87 and astrocyte HA1800 (fig S6), confirming the vital role of PLK1 on cell proliferation/survival. So PLK1 could be regarded as a marker to evaluate sensitive-insensitive effect of VAMP5 KD. Meanwhile compared to direct targeting PLK1, targeting VAMP5 using LV showed no influence on PLK1 expression and less harm on immortal astrocyte HA1800 viability (fig S6d-6e), which is foreseeable due to hardly expression of VAMP5 in astrocyte HA1800, indicating selectivity.

**Fig 4:**
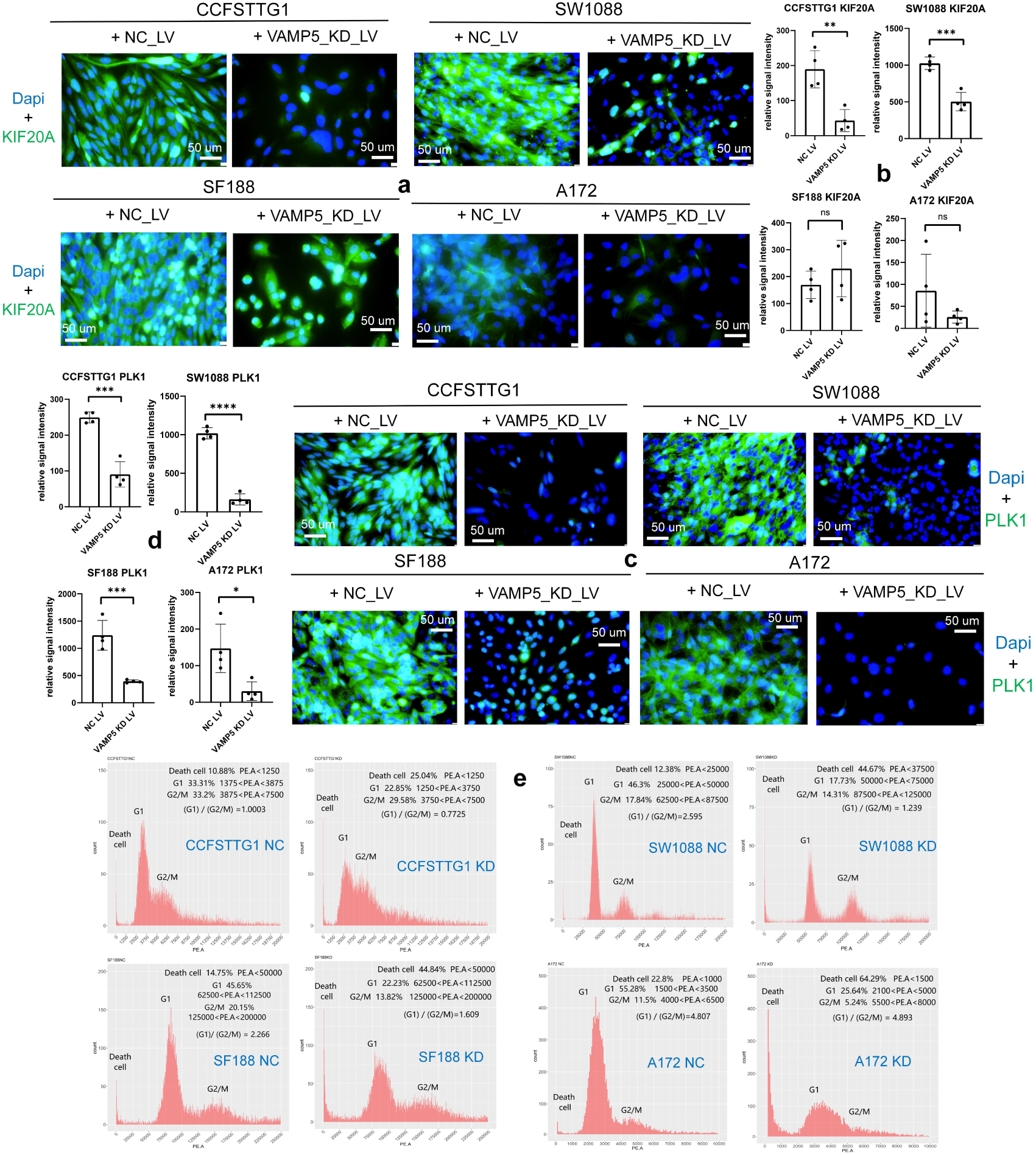
PLK1 were protein-level down-regulated in 4 sensitive cell lines with better consistency than KIF20A and caused cell cycle G1/G2M transition disorder; a: IF result staining KIF20A in which 4 parallel sights were calculated and one representative sight was exhibited; Same the situation in PLK1 results; b: semi-quantification results of KIF20A; c: IF results of PLK1; d: semi-quantifications of PLK1; e: Cell cycle testing results.

### MicroRNAs involved into mediation of VAMP5-KD induced downregulations of PLK1 and KIF20A

Tyramide Signal Amplification (TSA) multiple staining performed on in vivo tumor spheres sections to verify expressions and localizations of VAMP5 and PLK1 (or KIF20A). The results were beyond our expectations (fig 5a and S7) because VAMP5 was not apparently colocalized with PLK1 or KIF20A in NC or VAMP5 KD samples, while KIF20A and PLK1 generally showed better co-localization supporting the known conclusions [46]. The previous RNA-seq have shown clear RNA-protein inconsistency on PLK1 and KIF20A in SF188, and more inconsistency were discovered in other batches of VAMP5 KD or OE (fig 5b and S8). It seems that RNA-protein inconsistency on PLK1 or KIF20A is a very frequent situation. Additionally, GSEA on RNA-seq data of our cell line samples didn’t find enrichment of proteasome or ubiquitination related gene sets that will induce protein degradation without mRNA influence. Combined these 3 clues above, microRNAs may be possible intermediates because their imperfect complementary with 3UTR of targeted mRNA will silence translation but not always induce mRNAs degradation [51, 52], therefore tends to introduce mRNA-protein inconsistency. By microRNA-seq, 14 microRNAs in all were identified targeting PLK1 and KIF20A (fig 5c). CPM expression of 4 PLK1 targeting candidates were shown (fig 5d) and volcano plots of all 14 microRNAs plus CPM of 2 KIF20A targeting microRNAs with zero value (so not shown on volcano plot of corresponding cell lines) exhibited in fig S9. Mir-1301-3p looked like upregulating the PLK1 in 3 of 4 sensitive cell lines and mir-12135 compensated the rest unfitness of mir-1301-3p by upregulating PLK1 in A172 and meanwhile CCFSTTG1. Their pre-matures and predicted interaction structures with PLK1 were in fig 5e.

**Fig 5:**
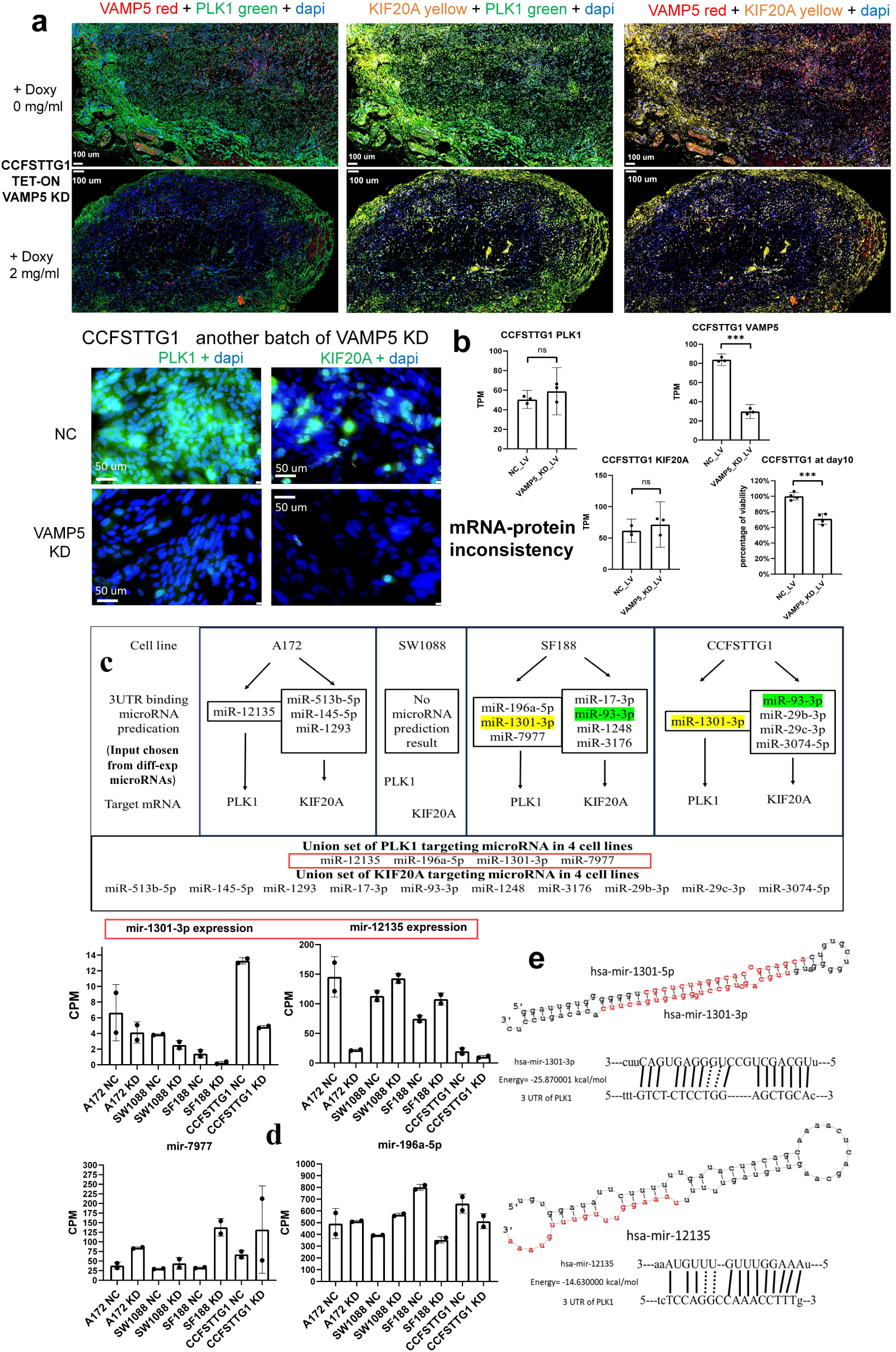
microRNAs, especially some up-regulation microRNAs may mediate VAMP5-KD induced downregulation of PLK1 and KIF20A; a: TSA triple staining result on in-vivo CCFSTTG1 tumor spheres indicating the expressions and localizations of 3 proteins; b: one representative example showing the possible mRNA-proteins inconsistency in cell line CCFSTTG1; c: All the microRNAs predicted targeting PLK1 and KIF20A in 4 cell lines; d: CPM barplots of 4 PLK1 targeting microRNAs; e: Pre-mature structures and predicted interactions between PLK1 3UTR and 2 representative up-regulation microRNAs, mir-1301-3p and mir-12135, respectively.

The up-reg microRNAs via interaction with 3UTR were first reported at 2022 [53] but compared to enormous amount of down-reg microRNAs, up-reg microRNAs were seldom. Therefore, we choose mir-1301 and mir-12135 for subsequent validations.

### Mir-1301-3p and mir-12135 were validated upregulating PLK1 in both sensitive and insensitive type of gliomas

Mir-12135 have not been reported before, while mir-1301-3p reported possessing both inhibition effect on breast cancer [54] and facilitation effect on prostate cancer stem cell [55], respectively, both via 3UTR interactions subsequent translational silence on different genes. Consequently, functional verification of these 2 microRNAs in glioma cells is exactly needed. All 4 VAMP5 KD sensitive cells plus one representative insensitive cell U87 were infected with same MOI of NC and sponge LV then tested cell viability, PLK1 protein-level expressions and sponge sequences using QPCR (designs overview in fig S10). Mir-1301-3p sponge showed common viabilities inhibition and PLK1 downregulation; Mir-12135 sponge showed same trends in A172, CCFSTTG1 and U87, viability inhibition without PLK1 downregulation in SW1088 and no inhibition effect at all in SF188 (fig 6). Hence, these 2 microRNAs are generally responsible for VAMP5-KD induced downregulation of PLK1 and growth inhibition simultaneously they themselves could also serve as independent treatment targets in both VAMP5-KD sensitive or insensitive type of gliomas. But individual differences should not be ignored just like behaviors of SF188 and SW1088 after mir-12135 sponge that one microRNA may play varying roles in different glioma cells.

**Fig 6:**
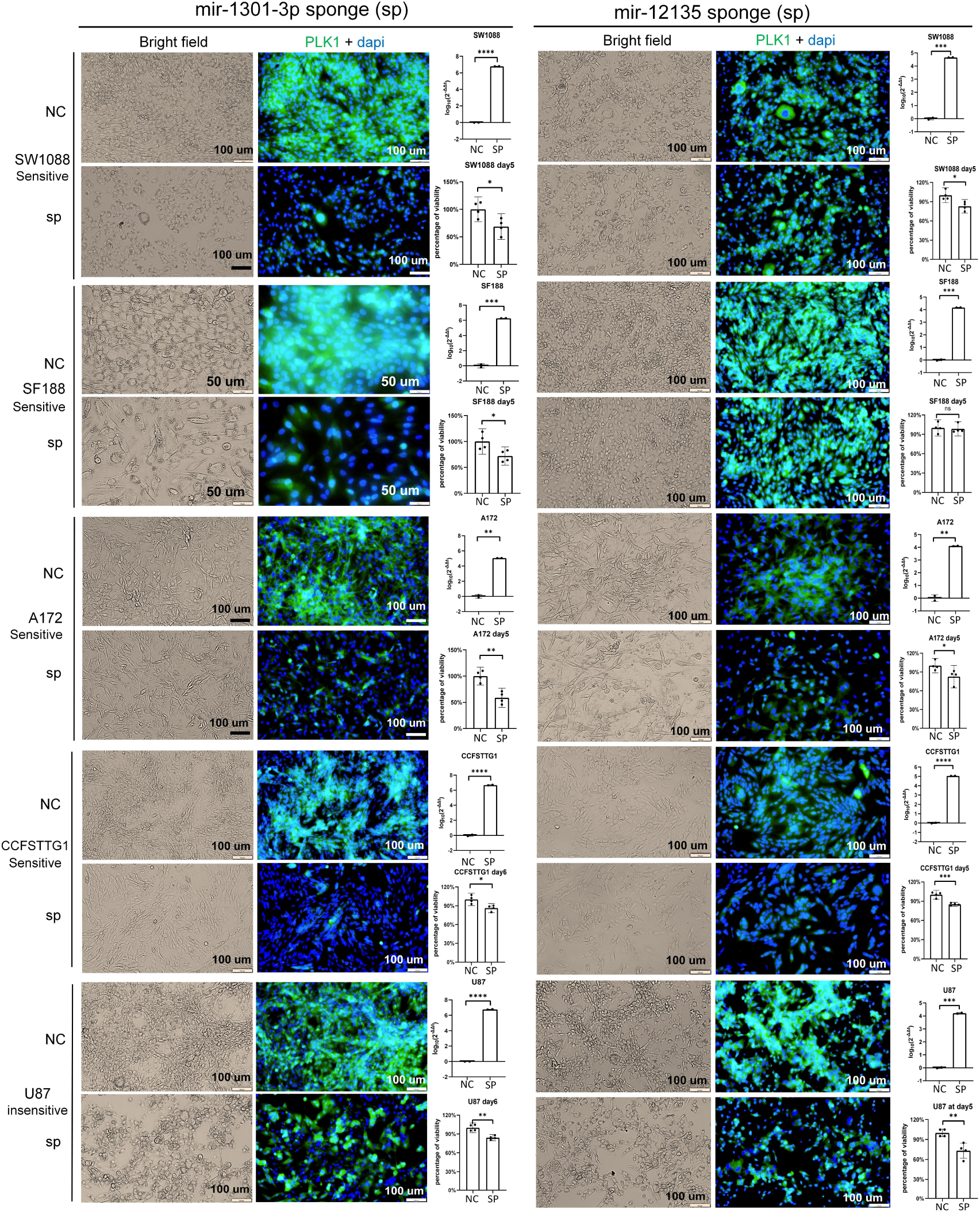
Blocking mir-1301-3p and mir-12135 using microRNA sponge method via lentivirus were validated overall effective for growth inhibition and PLK1 protein-level down-regulation in VAMP5-KD sensitive and insensitive cell lines, performed by IF, QPCR and viability test.

### NDRG4 is an RNA-level prediction marker distinguishing VAMP5-KD sensitive/insensitive type of gliomas as well as an enhancing target to convert insensitive type into sensitive type

Locating a prediction marker recognizing sensitive-insensitive gliomas is meaningful. We believe predictive information was included in RNA-seq data, so those from 4 sensitive cell lines and 3 insensitive cell lines were combined building training set, integratively using partial least square discrimination analysis (PLS-DA) and negative binomial test to search prediction marker and validate its prediction accuracy in new samples of testing set. NDRG4 was located as a suitable marker (fig 7a-c) in both training and testing set. The details of testing set results indicated that NDRG4 high-expressed LN229 and U251 showed significant viabilities inhibition and PLK1 protein-level decrease, while low NDRG4 expressed U87, MOGGUVW and LN308 (RNA/protein levels of VAMP5 not tested in this cell line) showed no VAMP5-KD induced alteration (fig S11). The testing results supported the predictions. FBS components influenced NDRG4 TPM expression and though not influencing the outcome of VAMP5 KD, using low-priced Chinese-originated FBS attenuated the discrimination effect of NDRG4 compared to high-priced Australia-originated FBS (fig 7d), which should be cautious during application.

**Fig 7:**
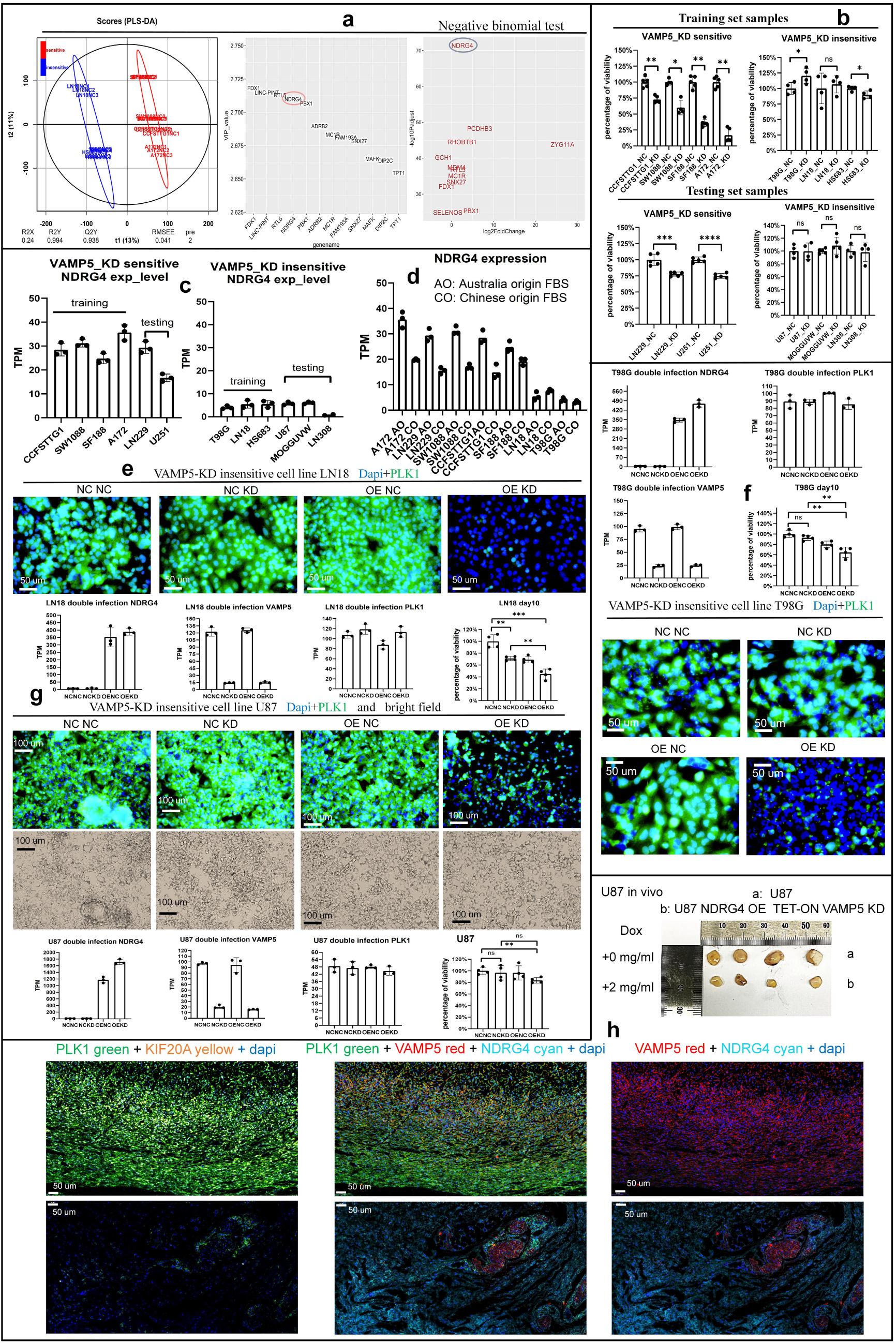
NDRG4 was identified as a prediction marker to distinguish VAMP5-KD sensitive and insensitive type of gliomas prior to real experiment. Furthermore, NDRG4 OE among VAMP5-KD insensitive cells could convert them into VAMP5-KD sensitive type; a: NDRG4 was located as the diff-expression marker between sensitive and insensitive types by integrative using PLSDA and negative binomial test; b: Both training samples and testing samples showed correct viabilities corresponding to sensitive and insensitive type, respectively; c: NDRG4 showed consistent high expression in 6 VAMP5-KD sensitive gliomas and comparatively consistent low expression among 6 insensitive type; d: Different FBS will influence expression of NDRG4, and Australia originated FBS showed more apparent distinguishing characteristic on NDRG4 than using Chinese originated FBS; e-h: NDRG4 OE in VAMP5-KD insensitive cell lines turned themselves into VAMP5-KD sensitive type that cell viabilities and protein-level expression of PLK1 were universally inhibited in: e: Cell line LN18 in vitro; f: Cell line T98G in vitro; g: Cell line U87 in vitro and h: Cell line U87 in vivo, using TSA multi-staining.

Inspired by the proven roles of PLK1 evaluation marker and NDRG4 prediction marker in gliomas VAMP5-KD system that possessing universal consistency, we wonder if NDRG4 OE fulfills the inhibition in VAMP5-KD insensitive type by converting the expression trend of PLK1. Experiments performed and results shown (fig 7e-h, fig S12). NDRG4 OE plus VAMP5 KD (OEKD) in insensitive type universally down-reg PLK1 protein-level expression apparently without RNA-level decrease and inhibited tumor growth both in vitro (fig 7e-g) and in vivo (fig 7h, fig S11). Cell cycle testing results showed that the PLK1 down-regulation induced by OEKD also altered the cell cycle states to various extents that differ from the situations of single VAMP5 KD among sensitive type (fig S13). Similarly to previously confirmed, KIF20A may also be reduced in part of cell lines via OEKD like U87 (fig 7h) and LN18 (fig S11b), but not true in all OEKD cells like T98G (data not shown), so KIF20A is not a suitable evaluation marker compared to PLK1.

### TNKS and VCAM1 were identified another 2 downregulated proteins induced by OEKD with consistency, in which loss of TNKS contributed to the reduction of PLK1 partly

The previous results have exhibited many mRNA-proteins inconsistencies, so we tried DIA-proteomics to explore other unknown influences of OEKD on insensitive type. The inaccurate quantifications of DIA-proteomics were observed (fig S14), so more technique should be applied to analysis proteomics data. The downstream analysis workflow shown as fig 8A, in which the vital principle is an experience that elements (proteins) within the same significantly enriched gene set are very possible to change synchronously following the global directions, maybe with various extents, so can be applied to exclude the doubtable quantifications (relies on luck partly). After performing the workflow, the down-regulated “RUIZ_TNC_TARGETS_UP” [56] in C2 set of MsigDb is the only enriched intersections between 2 OEKD cell lines meanwhile altered in same direction (fig 8b). The union set of core enrichment proteins were exhibited in heatmaps (fig 8c). Among these proteins, TNKS and VCAM1 were noticed very significantly lowered in one cell line, implying their actual reductions in others, respectively. TNKS supports the proliferation/progression of tumors via several accesses [57], and VCAM1 was discovered serving as immune checkpoint to help escaping from immuno-attack on tumors like hematopoietic cancer cells [58], both of which are beneficial to cancers. Therefore, multi-IF staining these 2 candidating proteins were performed on U87 and LN18 OEKD in-vivo tumor spheres, results shown that OEKD could induce their consistent reductions indeed (fig 8d-e). Followingly, we chose to validate the functions of TNKS, due to its influence on tumors themselves, by using its inhibitor XAV939 to mimic its down-regulation and demonstrated that function loss of TNKS inhibited tumors’ growth and caused the reduction of PLK1 on protein or RNA level (fig 8f-g), indicating its contribution on OEKD induced down-reg of PLK1 partly. Meanwhile, our experiments on another 2 VAMP5-KD sensitive cell lines, U251 and LN229 showed the possibility that disturbing TNKS also suppressed tumor growth through a PLK1 independent way (fig S15).

**Fig. 8:**
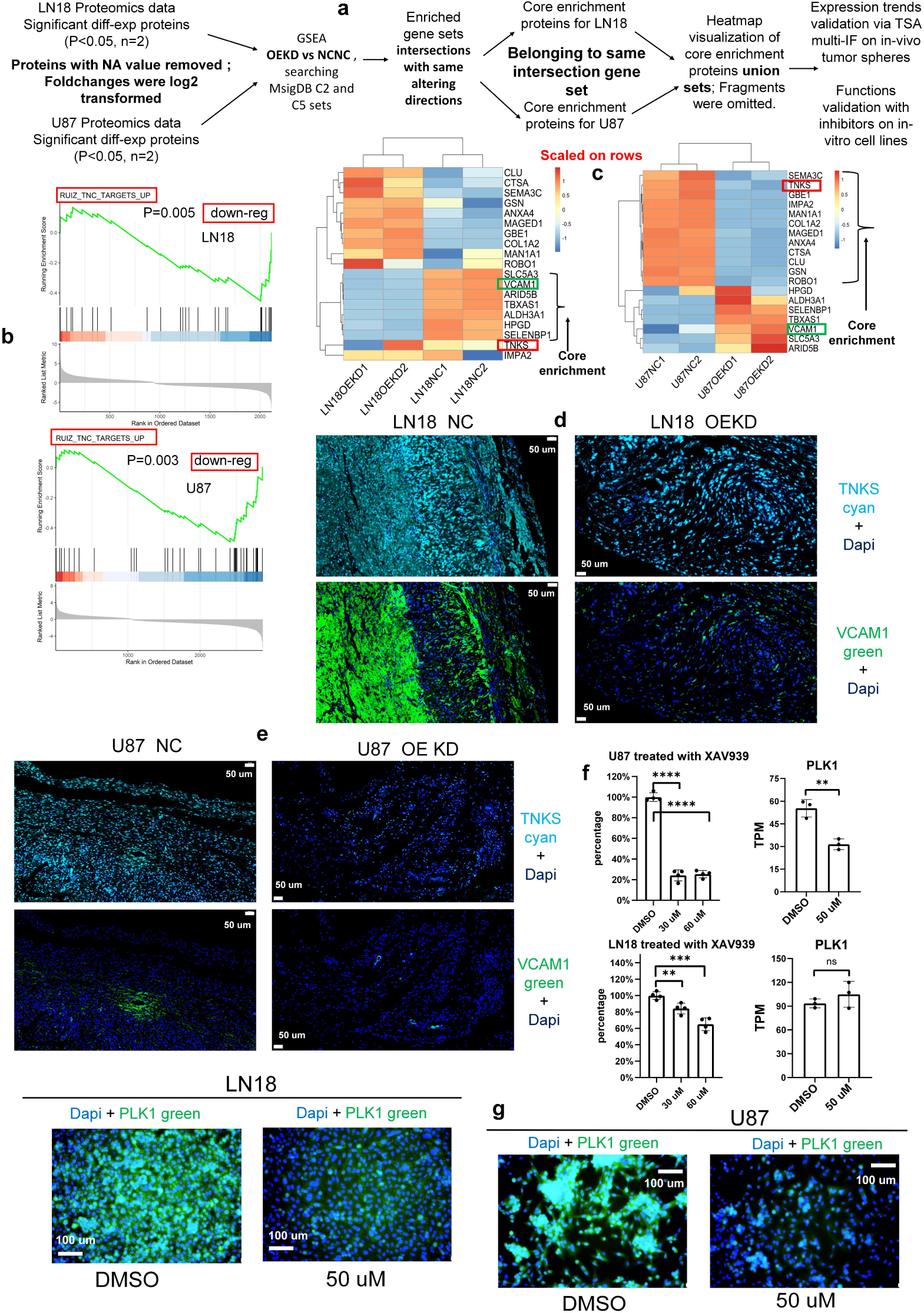
Applying proteomics-assisted exploration, TNKS and VCAM1 were found to be another 2 consistently down-regulated proteins induced by OEKD with specific supporting functions on tumors; a: Downstream data analysis pipeline to assure consistency between 2 cell lines and lower the misjudgment possibility due to inaccurate quantifications; b: Enrichment score plot of the consensus down-reg gene set intersection enriched in 2 cell lines; c: Heatmap of the union set of the core enrichment proteins in 2 cell lines belonging to the same down-reg gene set, in which TNKS and VCAM1 were noticed based on their alteration trends and background knowledge reviews; d: TSA multi-IF staining VCAM1 and TNKS on tumor spheres of cell line LN18 and e: U87; f: Cell viabilities and TPM RNA expression of PLK1 on cell lines U87 and LN18 treated with TNKS inhibitor XAV939; g: IF results staining PLK1 on 2 cell lines treated with TNKS inhibitor XAV939.

### VAMP5 KD may also induces correlated downregulation of membrane or cytoskeleton proteins, but except LGALS1, these correlations lack universal consistency

Based on the background knowledge reviewed initially, SNAREs participate in intracellular transportation and membrane proteins translocation/recycle. Thus, it’s natural to wonder if VAMP5 KD influences membrane proteins or other components. The RNA-seq results indicated that some membrane proteins and cytoskeleton components like FLNB that serve as a stabilizing linker between actin and inner tails of part membrane proteins [59, 60] may be downregulated by VAMP5 KD, but generally lack universal consistency among different cells (fig 9a-b), limiting the generalizability and application of this phenomenon. Simultaneously, among inconsistently-altered membrane proteins, CXCR4 and NTRK3 were reported their oncogenic roles [61, 62], so we also validated if they are effective targets independently or take effect in VAMP5-KD induced downregulated SW1088 cell using inhibitors in both altered SW1088 and nonchanged CCFSTTG1. Results showed that CXCR4 and NTRK3 were not responsible for glioma growth (fig S16), so the decrease of NTRK3 or CXCR4 in SW1088 was not the reason causing VAMP5-KD inhibition either. However, LGALS1 seems an exception that behaved universally downregulated in 5 of all 6 sensitive glioma cell lines plus one insensitive cell line and moreover, consistent with alteration trends among hundreds of TCGA samples (fig 9c, fig 1c). Protein-level verifications also demonstrated the generalizability of LGALS1-VAMP5 correlation and their co-localization trait (fig 9d). Hence, there is a potential to down-reg LGALS1, a classical and potent T cells/ NK cells inhibitor, via VAMP5 KD to attenuate tumor immune escape.

**Fig 9:**
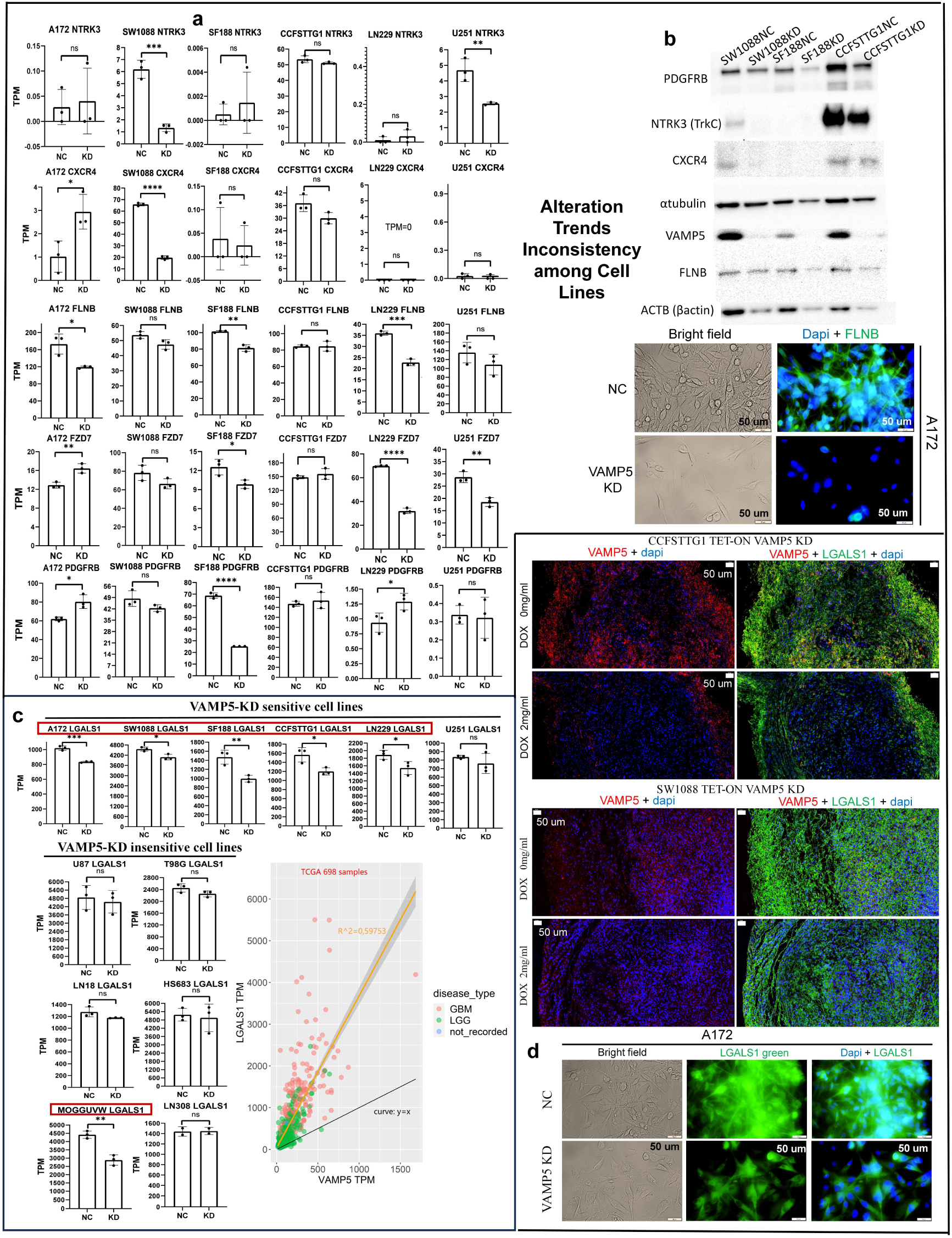
VAMP5-KD also induced down-regulation of some membrane proteins or cytoskeleton components, but except LGALS1, most of these correlations lack universal consistency among different cell lines; a-b: VAMP5-KD induced alteration inconsistency of membrane and cytoskeleton proteins on RNA level among VAMP5-KD sensitive cell lines and b: on protein level, tested via WB or IF; c: LGALS1 showed broader range of VAMP5 correlation among both cell lines and TCGA database samples on RNA level; D: LGALS1 was validated universally down-regulated on protein level following VAMP5-KD and colocalized with VAMP5 via TSA multiple staining of in vivo tumor spheres and in vitro single staining of cell line A172.

## Discussion

The original localization of VAMP5 as a possible target in gliomas relies on bioinformatic analysis of database samples. VAMP5 KD was invalid for growth inhibition in other type of cancers like rectal adenocarcinoma cells SW837, lung adenocarcinoma HCC827 and neuroblastoma SHSY5Y (data not shown), supporting our prediction to some extent. There are minor exceptions of gliomas that VAMP5 is not expressed on both RNA and protein levels like LN308 cell line (fig S11b), shows the diversity of gliomas and this type is certainly not suitable for VAMP5-targeting treatment.

In fact, the diversity of glioma cell lines response to same VAMP5-KD condition was shown all over this research such as the sensitive-insensitive response to VAMP5 KD. This distinction did not comply with official WHO grading on gliomas, because in sensitive cell lines, A172, SF188, LN229 belongs to grade IV GBM, SW1088, U251, CCFSTTG1 belong to lower-grade astrocytoma, while among insensitive cell lines, though U87, T98G, LN18 and LN308 belong to grade IV GBM, HS683 is oligodendroglioma and MOGGUVW is anaplastic astrocytoma, two patterns of classification not consistent. Consequently, the prediction role of NDRG4 (fig 7a-d) is a meaningful method to discriminate VAMP5-KD sensitivity of gliomas prior to real experiments that sensitive type possessing higher NDRG4 on RNA level than insensitive type. Furthermore, by NDRG4 OE among insensitive type, the subsequent VAMP5 KD will inhibit their growth via protein-level downregulation of PLK1, which turns them into VAMP5-KD sensitive type (fig 7E-H, fig S11). The existing studies on NDRG4 in gliomas showed conflicting results [63], including both up-regulation and down-regulation trends compared to brain normal tissues [64, 65], plus both facilitation and inhibition roles on proliferations of different gliomas [66, 67]. Our findings provided a new view on roles of NDRG4 in gliomas and applicable strategy for diagnosis and enhancing treatment. Certainly, experimental validations should be performed on more new samples to test the reliability. Additionally, the clear RNA-proteins inconsistency exhibited (fig 7e-g) suggested that microRNAs or other mechanism are possible to mediate NDRG4-OE-VAMP5-KD (OEKD) induced lowered-down of PLK1, further explorations needed.

Due to this mRNA-protein inconsistency, we also tried DIA proteomics to assist in finding other consistently-altered proteins unknown that induced by the OEKD above and validated via TSA multi-IF because of the possible inaccuracy on proteomic quantifications (fig S14). Fortunately, TNKS and VCAM1 were successfully located with consistent downregulation trends in OEKD of U87 and LN18 (fig 8a-e). TNKS is a promising target for cancer therapy due to its proliferation supporting roles on centromere elongation [57, 68], anaphase-metaphase transition of mitosis [57, 69], and positive regulation on canonical WNT pathway as well as transcriptional activator YAP1[57, 70–73], an important component in Hippos-YAP/TAZ pathway that is often activated abnormally in cancers; VCAM1 serves as an checkpoint responsible for immune tolerance in hematopoietic and leukaemic stem cells [58]. Therefore, parallel to PLK1, these 2 proteins may be another vital point responsible for OEKD inhibition. By independently inhibiting TNKS with inhibitor XAV939, the vital role of TNKS in gliomas was confirmed, and further validations showed that inhibition of TNKS may also downregulate PLK1 but with different down-reg extent or RNA expression trend (fig 8f-g), implied that there are more than one accesses mediating the reduction of PLK1 and growth inhibition of tumor cells in OEKD treatment on gliomas insensitive type, and the final effect were the net cumulative of these accesses. This hypothesis was also supported by experimental results of XAV939 treatment on 2 VAMP5-KD sensitive cell lines U251 and LN229 that inhibiting TNKS also suppressing tumor growth significantly without influencing PLK1 expression in some glioma cell lines (fig S15), demonstrated the existence of diverse suppressing mechanisms.

The different responses of sensitive cell lines to same VAMP5 KD like the different enrichment gene sets among 4 cells (fig 3a), also showed diversity. There is a point determining the repeatability of VAMP5-KD inhibition in CCFSTTG1 and SF188 that enough KD LV larger than threshold is necessary to overcome the cell resistance to VAMP5 KD (fig 2e) meanwhile several gene sets related to virus-infection response upregulated (fig 3a) [74, 75]. However, in SW1088 that did not exhibited similar gene sets upregulation, small amount of LV is enough for adequate VAMP5 KD. We have controlled same MOI of LV addition between NC/KD team to exclude nonspecific harm of viral particles so these up-reg gene sets may be responsible for KD resistance and need further investigation. The nonspecific harm of viral particles, however, should not be ignored. A172 behaved sensitivity to infection of NC LV, during which MOI of LV addition must be limited to lower than 20 to avoid nonspecific damage so that A172 did not show as apparent lower-down of VAMP5 than other 3 sensitive cell lines (fig S2, fig 3c). The addition amount of LV requires optimization for different glioma cell lines. If one glioma cell line is too sensitive to endure low level of NC LV addition (MOI<5), KD via viral method is not suitable for this cell because it’s not clear if the inhibition effect really originated from VAMP5 KD rather than nonspecific damage of viral particles. This situation was demonstrated true in astrocytoma cell lines SW1783 (data not shown).

The fourth aspect showing diversity is the alteration of some membrane proteins correlated with VAMP5 KD in parts of glioma cell lines like PDGFRB in SF188, FZD7 in LN229, U251, SF188 and CXCR4, NTRK3 in SW1088. This phenomenon also supported the current reviews on many SNAREs’ involvement in cancers about trafficking of membrane proteins. Inhibiting these SNAREs will block transportation towards correct membrane locations, followingly cause function loss and downregulation of influenced membrane proteins that are related to cancer’s progression or survival [30, 76]. But regarding VAMP5, first, its correlations with most of membrane proteins lack universal consistency among different glioma cell lines so the application was limited; Secondly, by inhibition of 2 representative non-universally altered proteins, NTRK3 and CXCR4 (fig 9a-b) on cell lines CCFSTTG1 and SW1088, they were validated independently as ineffective targets for gliomas inhibition (fig S16) so were not the reason causing tumor inhibitions even in VAMP5-KD altered cell line SW1088. The generalizability of VAMP5/membrane proteins correlations must be concerned. LGALS1, however, seems an exception possessing colocalization trait and comparatively wider correlation range with VAMP5 among both our cell lines and TCGA gliomas samples (fig 9c-d, fig 1c) and simultaneously, itself is a potent immune cells inhibitor [39], so it is possible to boost anti-tumor immunity by downregulating VAMP5 to lower LGALS1, and compared to direct targeting LGALS1, reducing the risk of potential self-immune attack problem [77, 78] , which also require further validations.

Similarly linking database analysis result, though we confirmed the universal consistency of protein-level PLK1 reduction in gliomas sensitive type and its constant role among gliomas insensitive type (fig 4c-d, fig S5), its correlation with VAMP5 among TCGA samples is very low (correlation coefficient lower than 0.1, data not shown). This may be due to the mediation of microRNAs introducing RNA-protein inconsistency so mRNA-seq analysis is not reliable to reflect real associations. The validated upregulation role of mir-1301-3p and mir-12135 on PLK1 was an unexpected discovery. Meanwhile IF results of HA1800 cell line treated with GSK461364A implied that PLK1 may not be very stable because in HA1800, though PLK1 exhibited low level of phosphorylation which should have been seldom influenced, the phospho-inhibitor of PLK1, GSK461364A [48] still induced real decrease of total PLK1 (fig S6a-b). Therefore, up-regulation microRNAs may be supplement strategy for PLK1. The direct blocking of mir-1301-3p or mir-12135 didn’t induce as apparent inhibition on viabilities as VAMP5 KD (fig 2b, fig 6). On the one hand, we only influenced one microRNA respectively compared to VAMP5-KD induced net cumulative of 14 in all so could not touch that extent (fig 5c-d, fig S9); On the other hand, maybe mediation of microRNAs are not the only reason responsible for PLK1/KIF20A downregulation simultaneously inclined to cause mRNA-protein inconsistency. The non-localization characteristic between VAMP5 and PLK1 or KIF20A in both NC and KD samples was another unexpected result (fig 5a and S7). Because the existing SNAREs researches supporting their interactions with other proteins usually relies on co-localization validations between SNAREs and related proteins in multiple staining assays, in which the correlated proteins are either located at other membrane structures mediating membrane fusions process by forming complexes with SNARE or served as cargos/co-components of the SNARE containing vesicles. Therefore, our result implied the VAMP5/PLK1 (or KIF20A) correlations not linked by 2 classical processes above directly, and the discovered microRNA intermediates explained this non-localization trait partly (not excluding other mechanisms). Finally, an interesting phenomenon was found that the inhibition on part of sensitive gliomas by VAMP5 KD seems non-relevant to classical death mechanisms like apoptosis or others (fig 3a, fig S4). We located KIF20A and subsequent PLK1 as vital points with efficiency based on this clue, but the reasons behind were to be investigated.

All in all, the overview of our discoveries in this research was summarized in fig 10. These systematic results provide new thoughts for gliomas targeting therapy with further development potentials.

**Fig 10:**
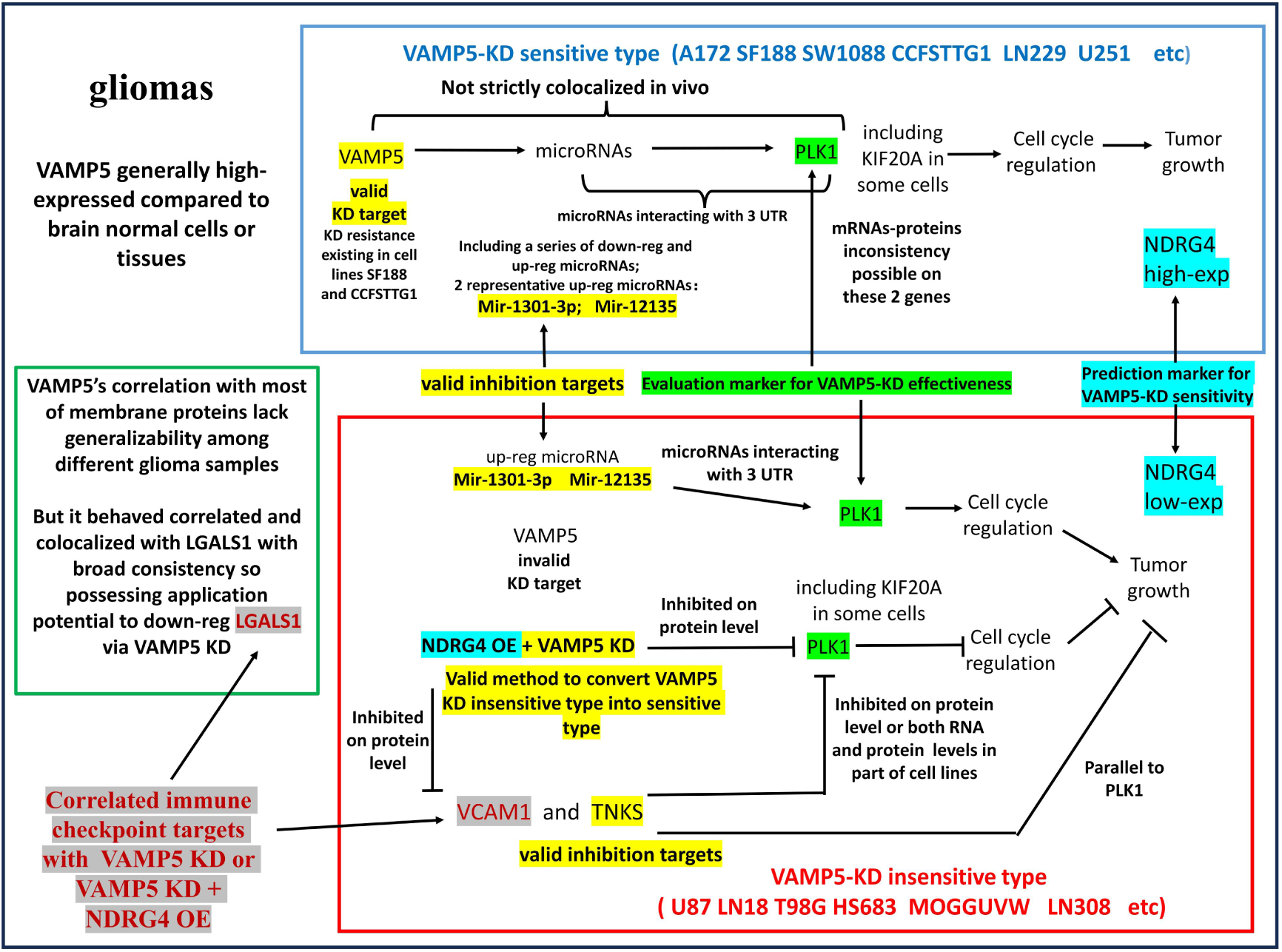
Overview of the new discoveries of this research.

## Material and method

### Database bioinformatic analysis

All the open-access read count RNA-seq data, and their corresponding clinical information downloaded from TCGA database grouping by primary site and disease type (table 1). Healthy samples downloaded from GTEX database. Read counts calculated into TPM using R language and Ensemble annotation. The non-overlapping exons base length computed by package “GenomicFeatures”. GSEA performed via “ClusterProfiler”, workflow summarized in fig 1a, and all elements of SNAREs gene set in table S1. Survival analysis performed through “survival” and “survminer” in which samples divided by expression median value. WGCNA was performed using package “WGCNA” with TPM as input to search the highly corelated genes in same module. TCGA samples and GTEX samples were combined, plotted by “ggplot2”; Gene track model was plotted by ggplot2 according to ensemble human genome annotations.

### Clinical sections and IHC

Paraffin embedded sections of glioma or paracancerous brain tissue from 8 patients were provided by neurosurgery department of Kaifeng Central Hospital. IHC assay operated following DAB staining protocol. Proteintech anti-VAMP5 (11822-1-AP) was primary antibody with a dilution of 1:100. Anti-VAMP5 primary antibody was replaced to BSA in NC (Negative control) team.

### Cell lines and cell culture

Glioma cell lines SW1088, CCFSTTG1 were purchased from MeisenCTCC, A172, MOGGUVW, SW1783 from CELLCOOK, T98G, HS683, LN229 from Hunan Fenghui Biotechnology Co., Ltd, LN18 from MINGZHOUBIO Co., Ltd, SF188 from IMMOCELL, U87, U251 from ServiceBio, LN308 from an anonymous laboratory. Immortal astrocyte HA1800 and HEB from JenioBio Co., Ltd. FBS from Australian (sigma F8687) and Chinese (TIANHANG 11011-8611) used respectively. Except CCFSTTG1, all the cell lines above cultured with DMEM + 10% FBS and 1% Penicillin-Streptomycin at 37℃ in 5% CO2 incubator. CCFSTTG1 cultured with RPMI1640 + 10%FBS +1% Penicillin-Streptomycin. The identities of cell lines were validated using STR analysis and were verified free of mycoplasma via PCR test.

### Lentivirus preparation and infection

Lentivirus packaged and concentrated via typical method and used for VAMP5 knocking down (KD), TET-ON KD, VAMP5-6×His overexpression (OE), NDRG4-6×His OE and microRNA mir-1301-3p and mir-12135 sponge. LV added into medium with varying MOI. In each batch of infections MOI of negative control (NC) and test team were equal. 1-5 ug/ml puromycin, depending on cell lines, added for 2 days for screening. Sequences cloned into viral vector plasmids shown in table S2.

### Colony formation assay and Giemsa staining

Cell suspension was diluted and counted by hemocytometer to add 3000 cells in 10cm diameter plates and incubated until colonies formed under microscope. Cells were fixed with 70% ethanol for 20 mins, stained with 1× Giemsa reagent (Beyotime C1033) for 30 mins then dried. Photos taken on the plate or under microscope.

### Western Blot

Anti-VAMP5 (ab216044,1:500), anti-α-tubulin (ab7291,1:2500), anti-β-actin (ab8226,1:1000), anti-FLNB (ab224334, 1:1000), anti-NTRK3 (ab240651, 1:1000), anti-CXCR4 (ab181020 1:1000), anti-PDGFRB (proteintech 13449-1-AP, 1:1000), donkey-anti-rabbit HRP secondary antibody (proteintech, SA00001-9, 1: 1000), and donkey-anti-mouse HRP secondary antibody (proteintech,SA00001-8, 1: 1000) were used.

### Cell apoptosis test

GreenNuc^TM^ Caspase-3 Assay Kit (Beyotime, C1168M) applied to test apoptosis based on caspase-3/7 activity. Apoptosis level evaluated by the numbers of green stained cell nucleus. Camptothecin (Beyotime, ST2123) dissolved in DMSO and added to cell at the final concentration of 10μM, preserved for 12h to induce apoptosis as positive control.

### Cell viability assay

CCK8 reagent used for cell viability assay. Old medium removed, and new medium mixing 1%(v/v) CCK8 added to cell. Same volume of CCK8 mixed medium was added to cell-free wells as blank control. After cultured in incubator at 37 ℃ for 70 mins, the CCK8-mixed medium was equal-volume-added to 96-well plate to test OD450 on microplate reader. 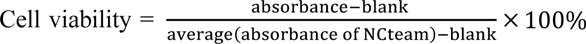.

### Immune Fluorescence and semi-quantification of target protein

Following regular method. 4% formaldehyde/PBS at 4℃ for fixation, 0.2% TritonX-100/PBS at room temperature for permeabilization. 10% goat serum (Solarbio SL038) dissolved in BSA for blocking and antibodies dilution. Anti-VAMP5 (sigma HPA035082, 1:100), anti-KIF20A (proteintech 15911-1-AP, 1:200), anti-PLK1 (abcam ab189139, 1:250), anti-phospho-PLK1(MCE HY-P80474, 1:50), anti-FLNB (abcam, ab224334, 1:100), anti-LGALS1(abcam ab108389, 1:100), anti-6×His tag (CST #12698, 1:250) were primary antibodies, and goat-anti-rabbit 488 (abcam ab150077, 1:500) was secondary antibody.

Samples between control team and test team within same batch have equal exposure time and same background removal. ImageJ applied to semi-quantify target protein in one picture following the equation *r*elative signal intensity 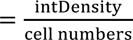. 4 parallel pictures analyzed and one representative picture in control and test team exhibited, respectively.

### RNA-seq experiment and data analysis

RNA extraction and library construction via protocols. Sequencing performed on NovaSeq 6000 platform. Software “fastp”, “hisat2”, “samtools”, “stringtie” accompanied by “prepDE” python script used in workflow. Ensemble annotation for reference annotation. R Package “DESeq2” used to perform diff-exp analysis, followingly “clusterProfiler” for GSEA analysis, C2 and C5 sets in Molecular Signature Database (MsigDB) were library searched. “rtracklayer” for TPM value extraction and arrangement, “ggplot2” and “pheatmap” for GSEA results and expression heatmap visualizations.

### Drug treatment to cell

GSK461364A (MCE, HY-50877), doxycycline hydrochloride (MCE, HY-N0565A), XAV939 (MCE, HY-15147), Larotrectinib (Aladdin, L172696), Plerixafor (Aladdin, P128026), Temozolomide (Macklin, T838219) applied to treat cell with corresponding concentrations, dissolving reagents and treating time.

### Cell cycle test

Cell Cycle and Apoptosis Analysis Kit (Beyotime, C1052) applied for cell cycle test. At least 4×10^5^ cells digested by 0.25% trypsin-EDTA, filtered by 40 micrometer cell strainer, centrifugated at 2000g, re-suspended by PBS and slowly added into pre-cooled ethanol. The final concentration of ethanol was 70%. Cell suspensions stored at -20℃ for 24h for fixation and permeabilization. Fixed cells centrifugated at 1500g and resuspend with propidium solution, stained avoiding light at 37℃ for 30mins, finally performed analysis on flowcytometry. The output data browsed and visualized in cell count histogram using R package “flowCore” and “ggplot2”.

### Animal experiments and TSA staining

5×10^6 TET-ON VAMP5-KD stable cell lines, NDRG4 OE TET-ON VAMP5-KD stable cell lines, or negative control cell lines including CCFSTTG1, SW1088, U87 and LN18 digested, mixed with 50%-medium-diluted matrix-gel (GelNest 211252), injected into left flank of nude mice subcutaneously. 2 mg/ml doxycycline chloride supplied in feeding water since day 3. Mice executed at day 17 and photos of removed tumors taken. Paraffin sections preparation of tumors and Tyramide signal amplification (TSA) multi-IF proteins staining performed as protocols. Anti-VAMP5 (sigma-HPA035082, 1:100), anti-PLK1 (CST-208G4, 1:200), anti-KIF20A (proteintech 15911-1-AP, 1:2000), anti-LGALS1 (abcam ab108389, 1:500), anti-NDRG4 (proteintech 12184-1-AP, 1:200), anti-TNKS (proteintech 18030-1-AP, 1:3000), anti-VCAM1 (Servicebio GB113498, 1:2000) were primary antibodies used.

### MicroRNA-seq and data analysis

Sequencing performed on NovaSeq 6000 following protocols. Reference genome index built from GRCH38 human genome by software “Bowtie2”. “miRDeep2” for reads mapping, quantification and pre-mature structures prediction. Known human microRNA mature and pre-mature sequences from miRBase were annotations. DESeq2 used for diff-exp analysis that microRNA with |log2FoldChange|>1 and P-value<0.05 considered as diff-exp candidates. Software “bedtools” and “miRanda” for 3 UTR sequences extraction from GRCH38 and targeted microRNAs predictions based on diff-exp candidating microRNAs.

### Real-time quantitative PCR

Real-time quantitative PCR (QPCR) used to test artificial sponge sequences blocking mir-1301-3p and mir-12135, respectively, primers sequences in table S3, covering locations and design overview in fig S10. Experiments performed following typical protocols. β-actin was housekeeping genes.

### Sensitive-insensitive prediction marker identification

Training set RNA-seq data analyzed via packages “DESeq2” and “ropls”, using negative binomial test and PLS-DA respectively, locating marker genes by intersection between 2 analysis results and expression trends validation in RNA-seq testing set of new samples. Read counts were input of DESeq2 and TPMs were that of ropls.

### Proteomics and data analysis

DIA (Data independent acquisition) quantitative proteomics applied to assist searching altered proteins on U87 and LN18 cell lines imposed to VAMP5 KD plus NDRG4 OE condition in vitro (n=2). Pre-treatment of samples before injections followed typical protocols. Vanquish Neo UHPLC (Thermal Scientific) and Orbitrap Astral (Thermal Scientific) platforms were used instruments; DIA-NN (version 1.8) software for raw MS (Mass spectrometry) data processing and proteins identification/quantification via DIA algorithm, with Uniprotkb-Homo Sapiens as reference database and Qvalues<0.01 as threshold. The output signal intensities of corresponding proteins were input data of downstream analysis including GSEA (via R package clusterProfiler), hierarchical cluster heatmap visualizations (via package pheatmap) and fundamental barplot visualizations (via Prism Graphpad), details shown as fig 8A and fig S14.

### Statistics and graphic plot

T-test and negative binomial test were used. Prism Graphpad, ggplot2 and other R packages mentioned before for results visualization.

### Data availability

Raw data will be supplied by first author for reasonable request.

## Supporting information

all supplemental figures

all supplemental tables

## Acknowledgements

Not applicable.

## Authors’ contribution

Yinghao Zhang: Conceptualization, funding acquisition, data curation, formal analysis and investigation; Yisheng Liu: Funding acquisition, resources and project administration; Wanhong Zhang: Resources; Haoran Wang: Formal analysis and investigation; Jinhai Wang: Investigation and validation; Yufang Sui: Validation and visualization.

## Funds

Not applicable.

## Ethics approval

Animal experiment was performed in accordance with guidelines and regulations (reference numbers: HUSOM2025-014) approved by Committee of Medical Ethics and Welfare for Experimental Animals, Henan University School of Medicine. Application of human samples was approved by Kaifeng Central Hospital Medical and Scientific Research Ethics Committee (Ethical Committee number: 2021LL012-KFZXYY). All the participants consented to participate in the study.

