## Supplementary material for "VAMP5 is a novel target for selective inhibition of gliomas with high NDRG4 expression via downregulating PLK1 and beyond": all supplemental figures

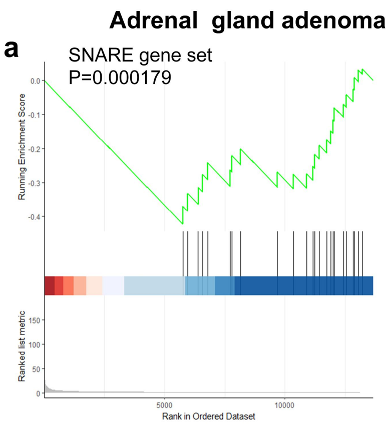


| genename | meanTPMfoldchange | pvalue |
|---|---|---|
| STX5 | 1.787045 | 1.25E-05 |
| STX6 | 1.742514 | 8.40E-06 |
| YKT6 | 1.727859 | 5.03E-05 |
| STX18 | 1.708761 | 6.11E-05 |
| STX17 | 1.62911 | 0.00169 |
| SNAP29 | 1.62231 | 4.93E-05 |
| STX10 | 1.597317 | 6.83E-07 |
| GOSR1 | 1.498424 | 9.20E-05 |
| VAMP2 | 1.457703 | 0.002944 |
| VTI1A | 1.4237 | 0.001476 |
| BET1L | 1.405047 | 0.0017 |
| BNIP1 | 1.399997 | 0.027542 |
| SEC22B | 1.387706 | 0.020227 |
| GOSR2 | 1.362455 | 0.005166 |
| VTI1B | 1.347139 | 0.02287 |
| SNAP47 | 1.341174 | 0.015361 |
| STX12 | 1.33617 | 0.034875 |
| VAMP4 | 1.30239 | 0.01765 |
| VAMP3 | 1.290466 | 0.015425 |
| STX16 | 1.256856 | 0.006667 |
| USE1 | 1.251321 | 0.028305 |
| STXBP3 | 1.204421 | 0.045965 |
| PRKCA | 0.516521 | 0.032716 |

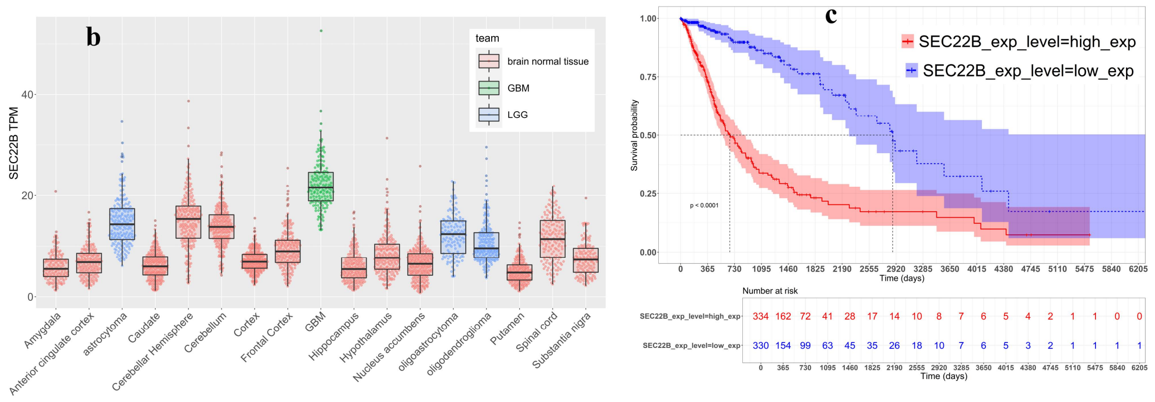


**Fig S1: additional information on database analysis**; a: Enrichment score plot and corresponding core enrichment genes in another cancer type SNAREs were significantly enriched: Adrenal gland adenoma & adenocarcinoma; b: Box-beeswarm plot of SEC22B expression among TCGA glioma samples and GTEX normal samples; c: Survival analysis result of influence of SEC22B expression level on TCGA glioma samples.

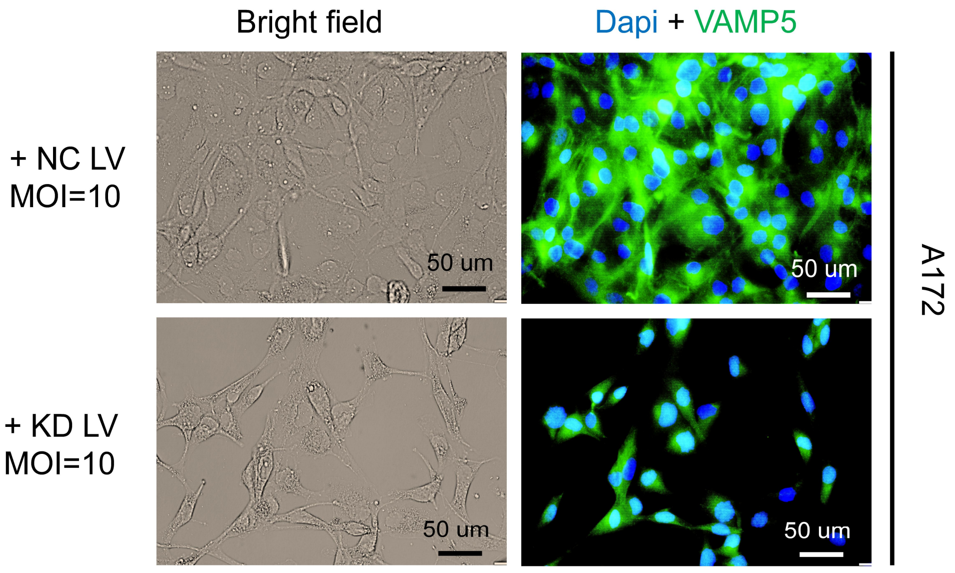


Fig S2: **IF result staining VAMP5 on A172 cell line.**

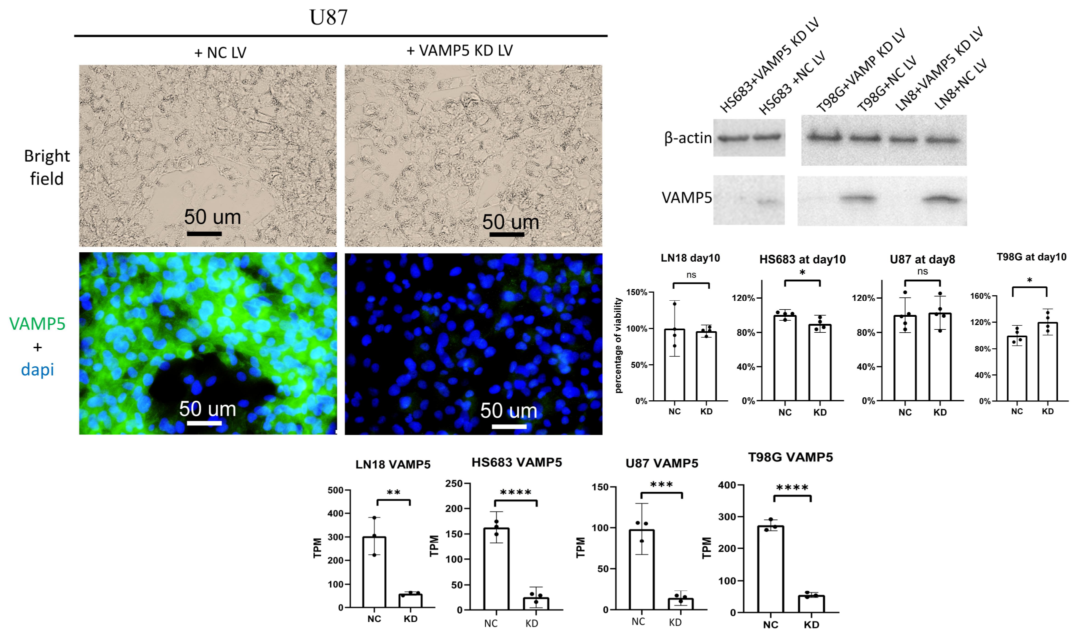


Fig S3: **Cell viabilities and VAMP5 expression in VAMP5-KD insensitive type of cell lines**

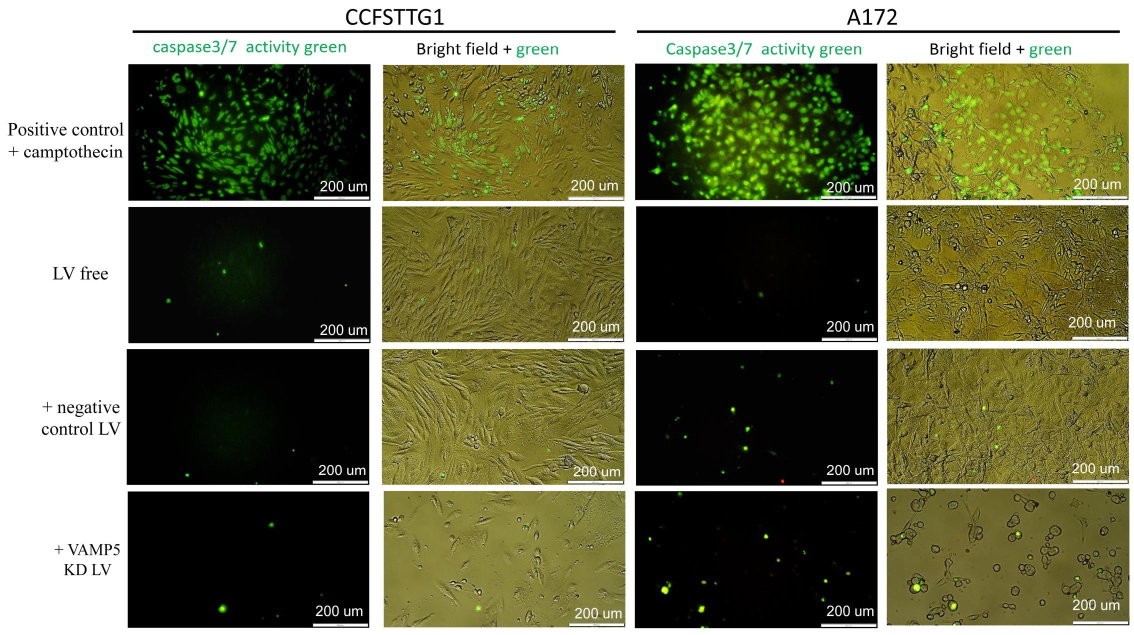


Fig S4: **VAMP5-KD induced growth inhibition of gliomas was not fulfilled through apoptosis.**

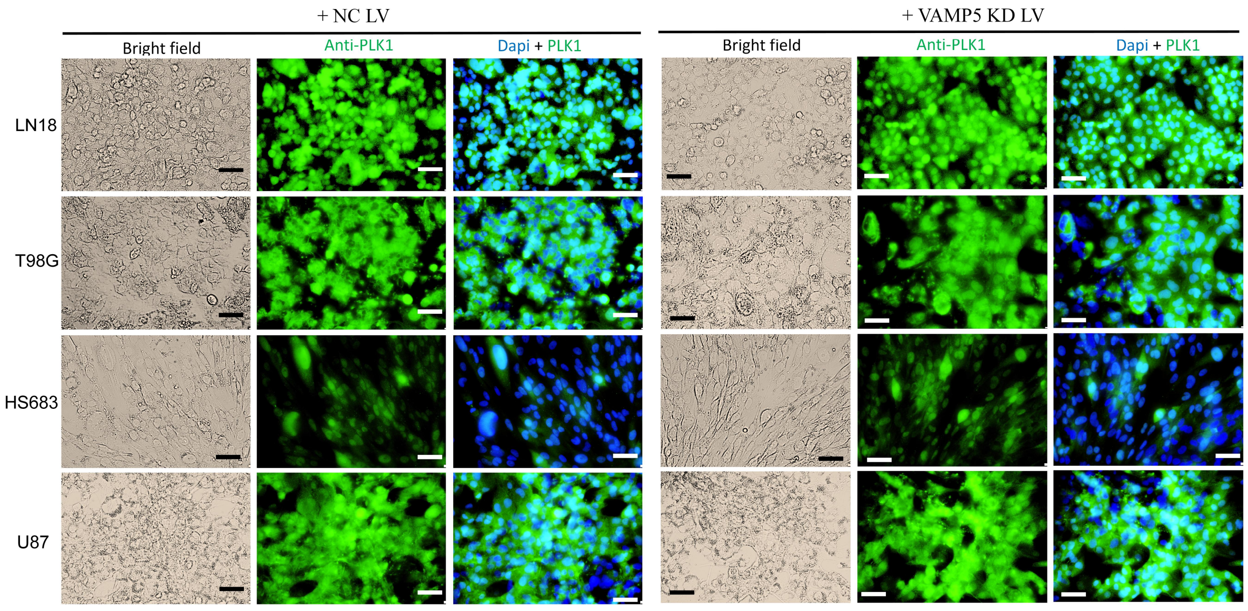


Fig S5: **Compared to gliomas of VAMP5-KD sensitive type that KD VAMP5 will cause universal downregulation of PLK1, KD VAMP5 among insensitive type of gliomas did not induce alteration on PLK1 consistently**. Scales stand for 50um in length.

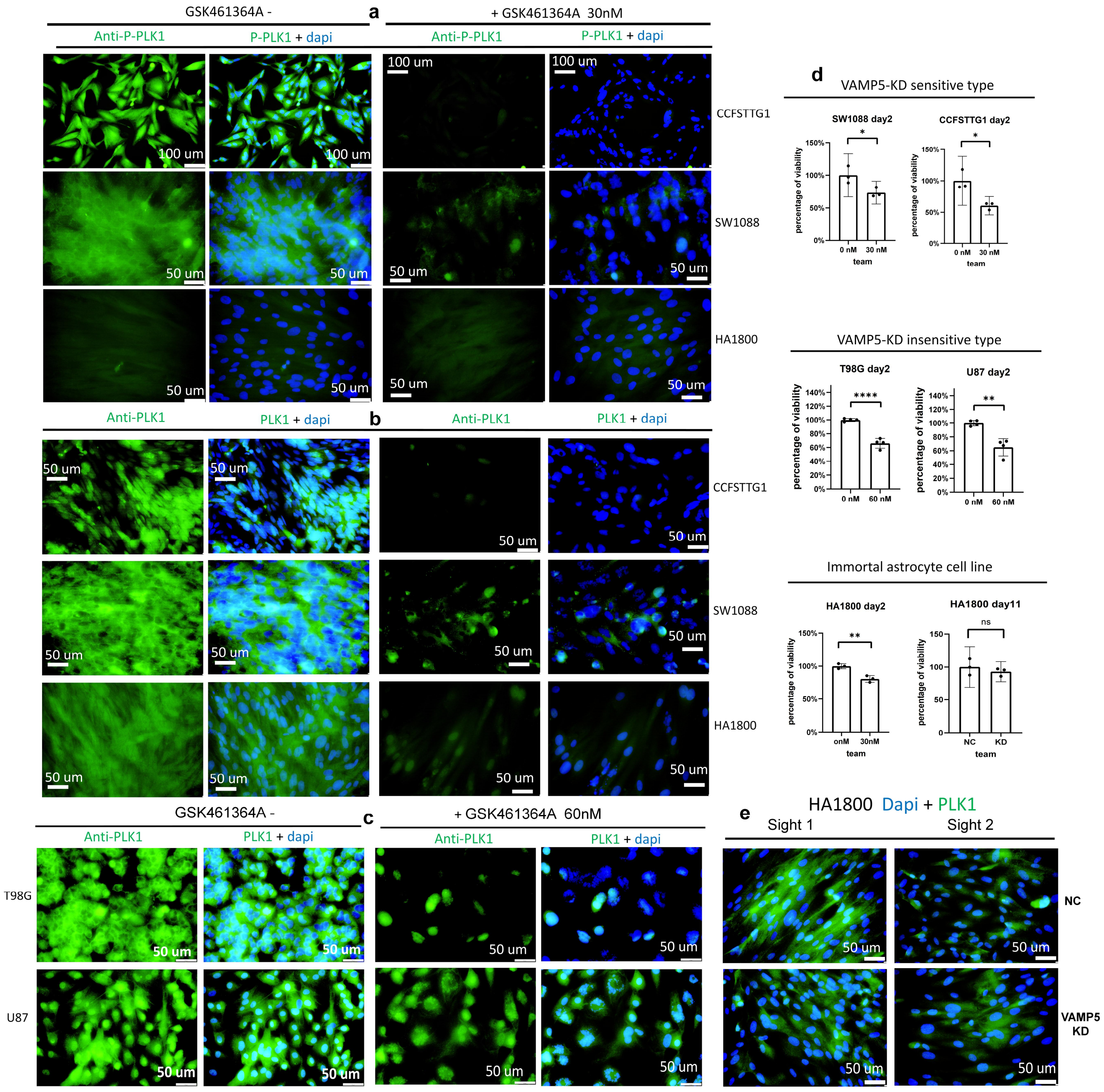


Fig S6: **PLK1 is vital for growth/survival of gliomas belonging to both VAMP5-KD sensitive and insensitive type as well as normal astrocyte cells, and compared to directly targeting PLK1, aiming at VAMP5 showed less harm on normal cells**; a: IF results staining P-PLK1 and b （staining PLK1） on VAMP5-KD sensitive cells or immortal astrocyte cells HA1800 treated with PLK1 inhibitor GSK461364A; c: IF staining PLK1 on VAMP5-KD insensitive cells treated with GSK461364A; d: Cell viabilities of GSK treated cells and HA1800 adding NC or VAMP5 KD LV; e: IF staining PLK1 on HA1800 adding NC or VAMP5 KD LV.

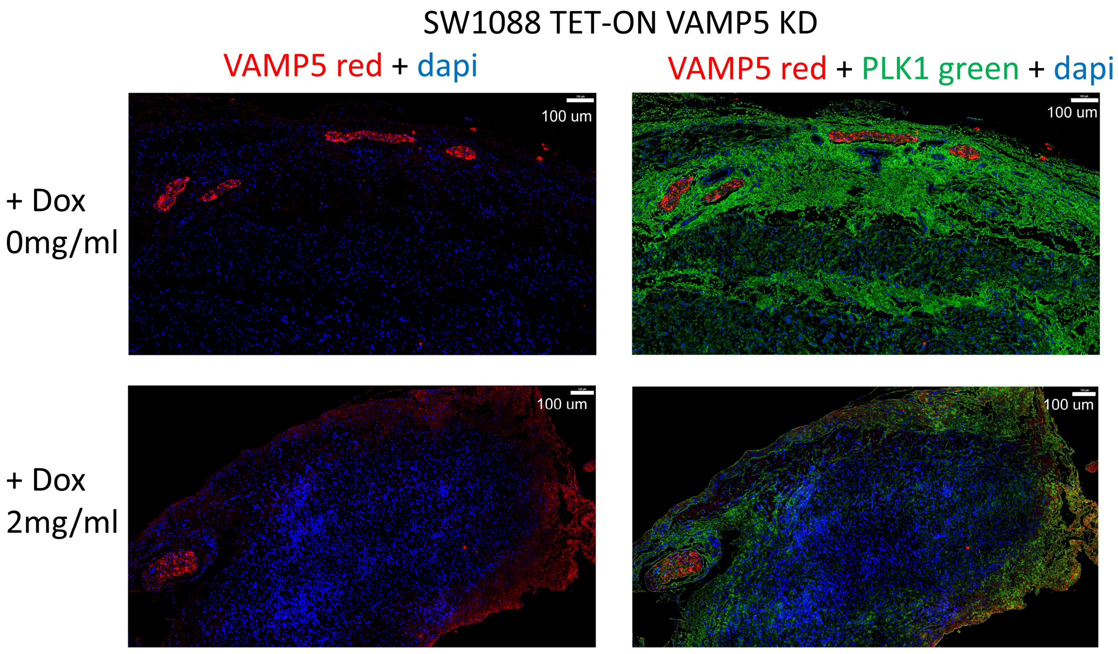


Fig. S7: **TSA double staining VAMP5 and PLK1 on SW1088 tumor spheres removed from xenograft model, in which PLK1 was downregulated and non-colocalized with VAMP5.**

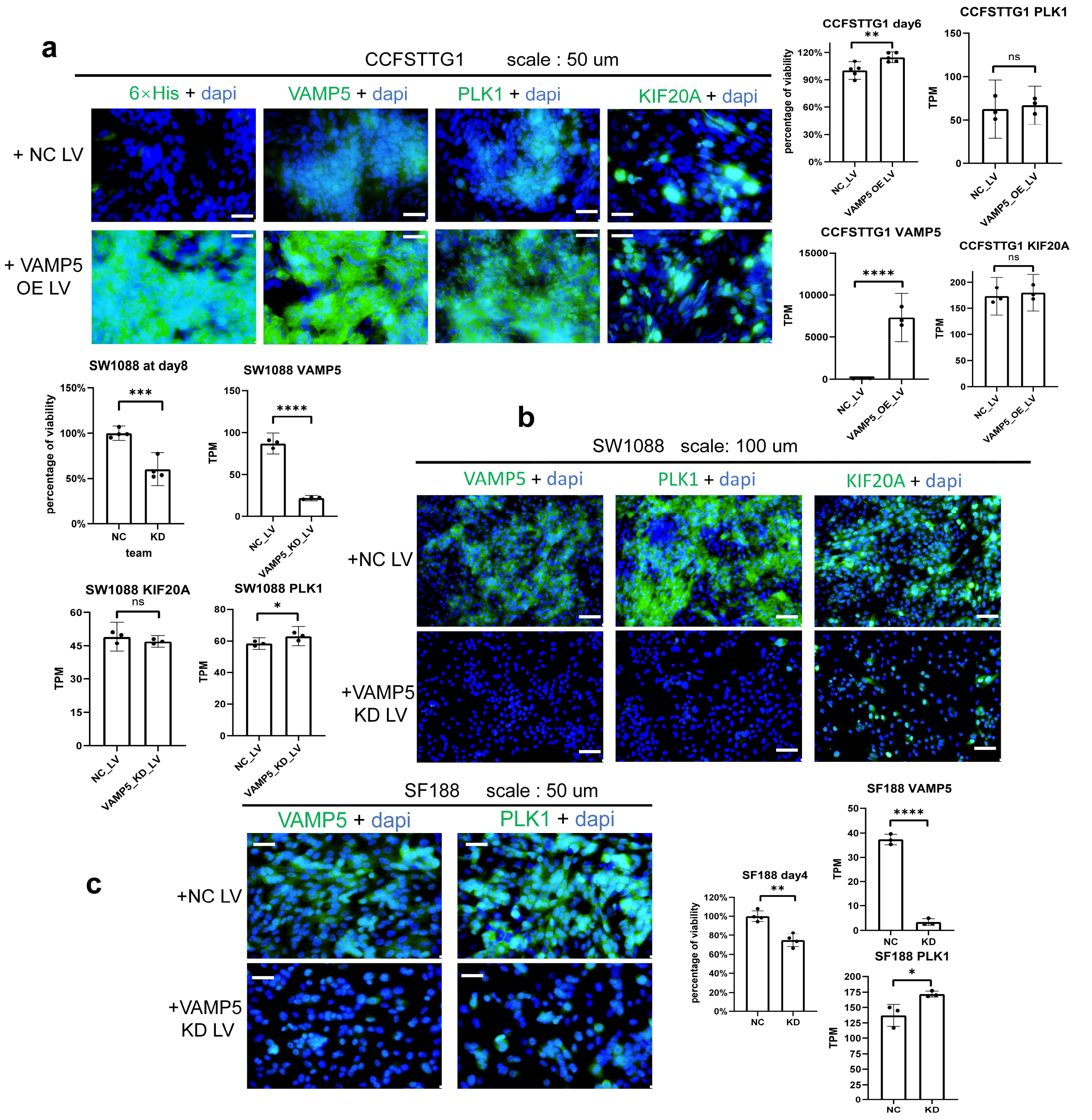


Fig S8: **There exists RNA-proteins inconsistency in other batches of assay performed on A: CCFSTTG1 VAMP5 overexpression; B: SW1088 VAMP5 KD and C: SF188 VAMP5 KD**.

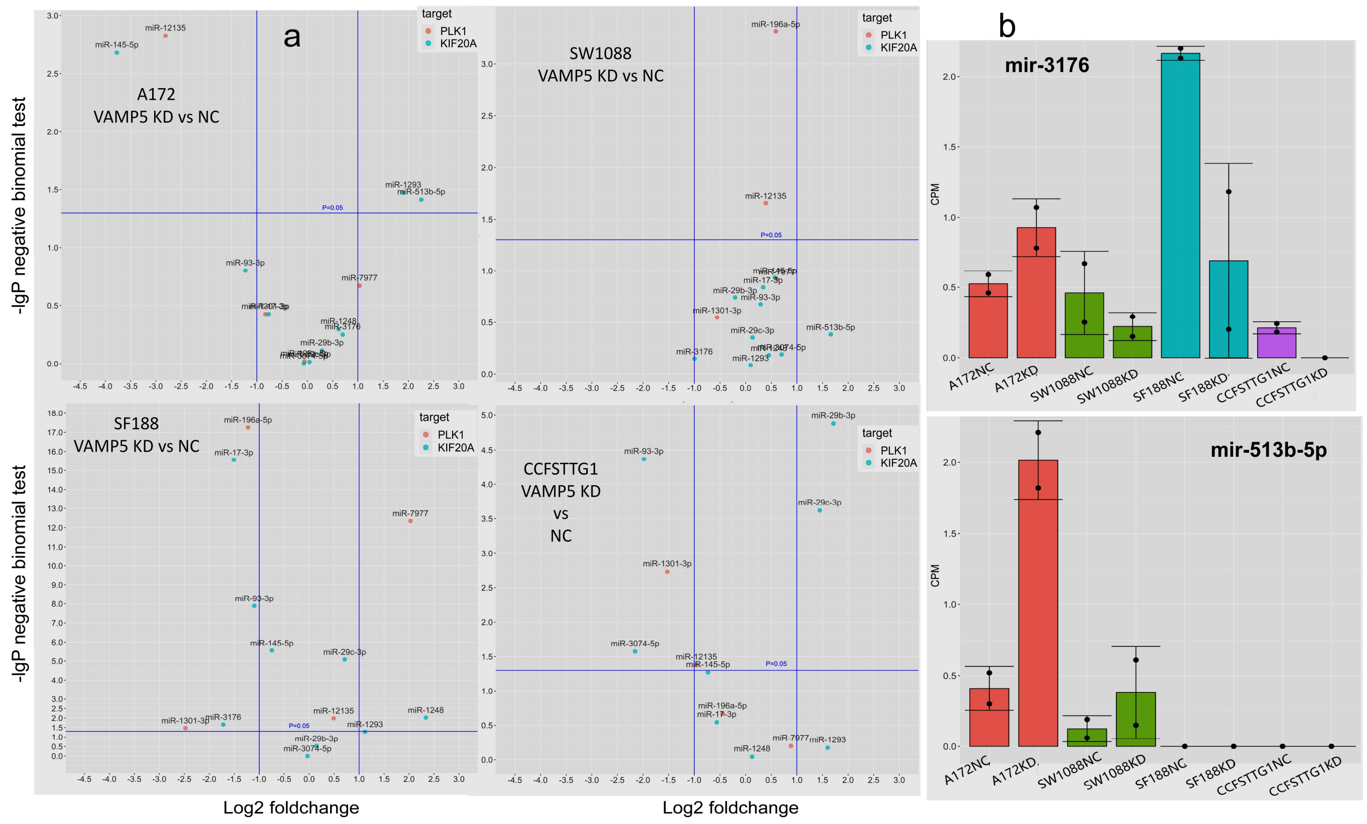


Figure S9: **Additional results on microRNA-seq of 4 VAMP5-KD sensitive cells;** a: Volcano plots of diff-exp microRNAs with predicted targets in corresponding cell lines; b: CPM barplots of 2 KIF20A targeting microRNAs that existed zero values so unable to be shown on volcano plot of corresponding cell lines.

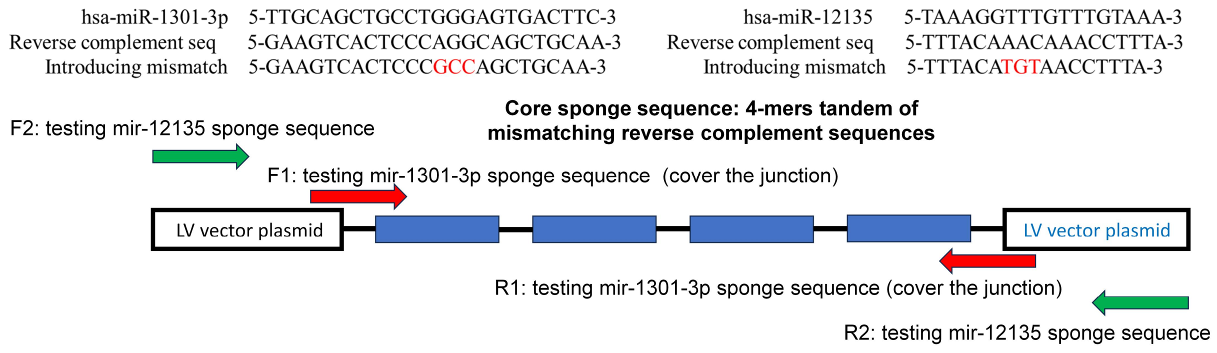


Fig S10: **Design of microRNA sponge sequences and their corresponding testing QPCR primers.**

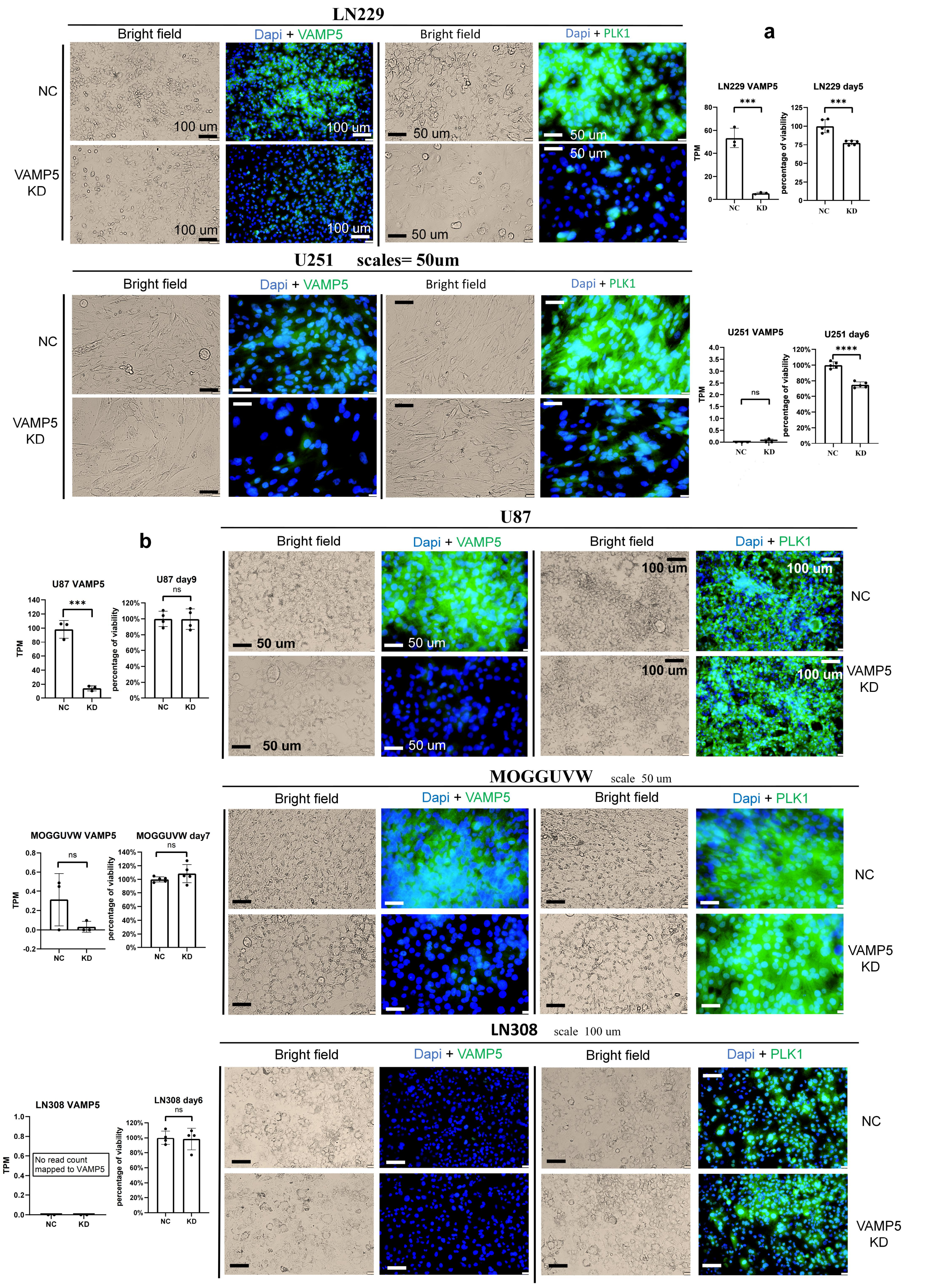


Fig S11: **Experimental validation results on prediction of VAMP5 KD sensitivity in testing set samples;** a: Experiment results of 2 predicted belonging to sensitive type, LN229 and U251; b: Results of 3 predicted insensitive type of cell lines, U87, MOGGUVW and LN308.

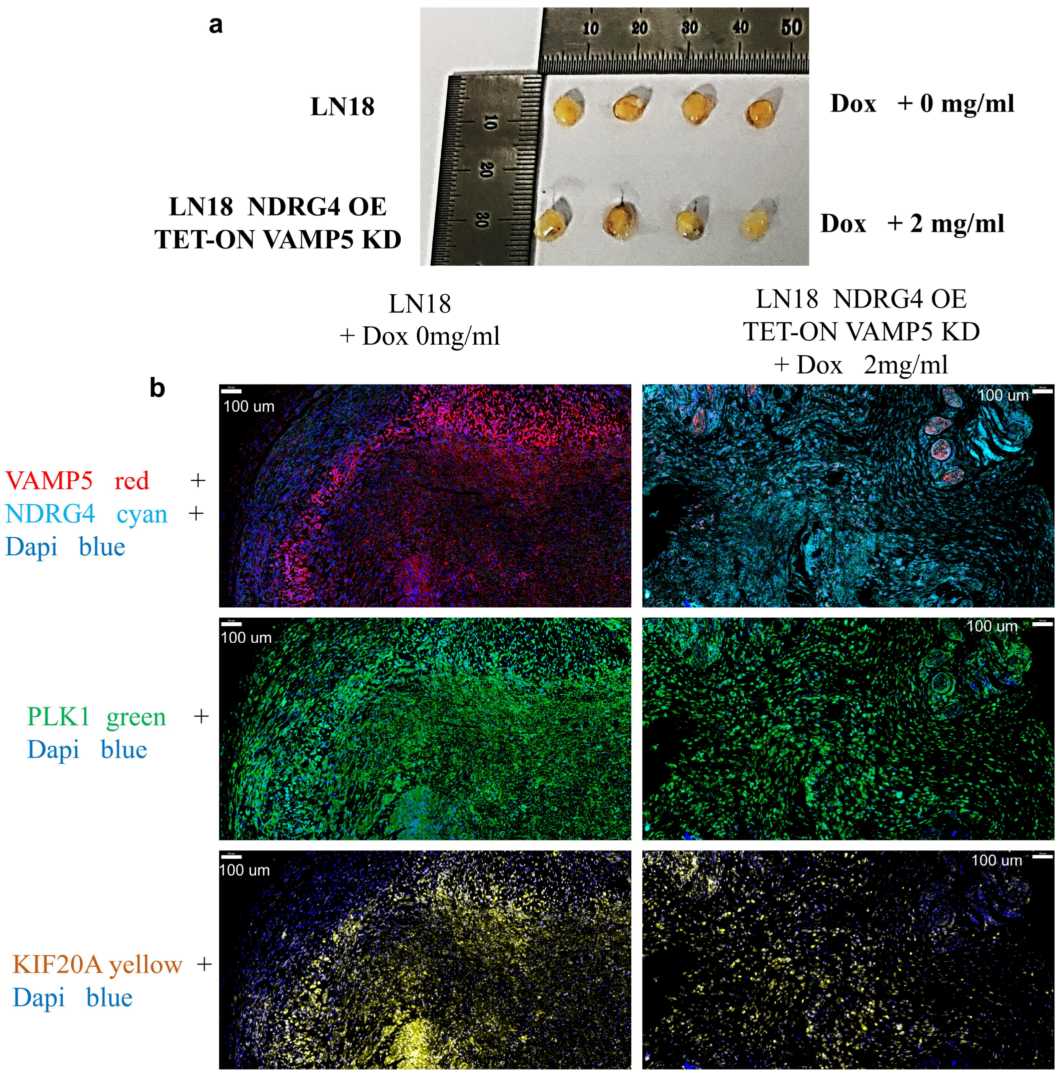


Fig. S12: **Results of VAMP5 KD plus NDRG4 OE on cell line LN18 in vivo**; a: Tumor spheres sizes comparation, in which the samples in NC team were more compact than OEKD; b: TSA staining of VAMP5, NDRG4, PLK1 and KIF20A on LN18 tumor spheres.

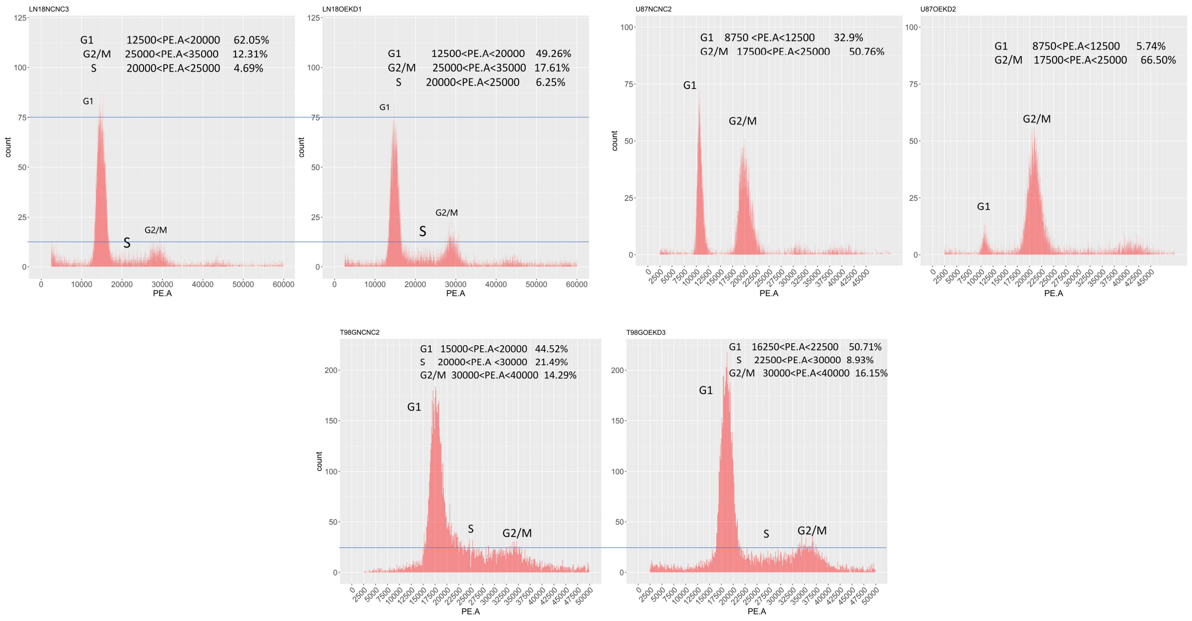

Fig. S13: **Cell cycle testing result of NDRG4 OE + VAMP5 KD on insensitive type of glioma cell lines LN18, U87 and T98G.**

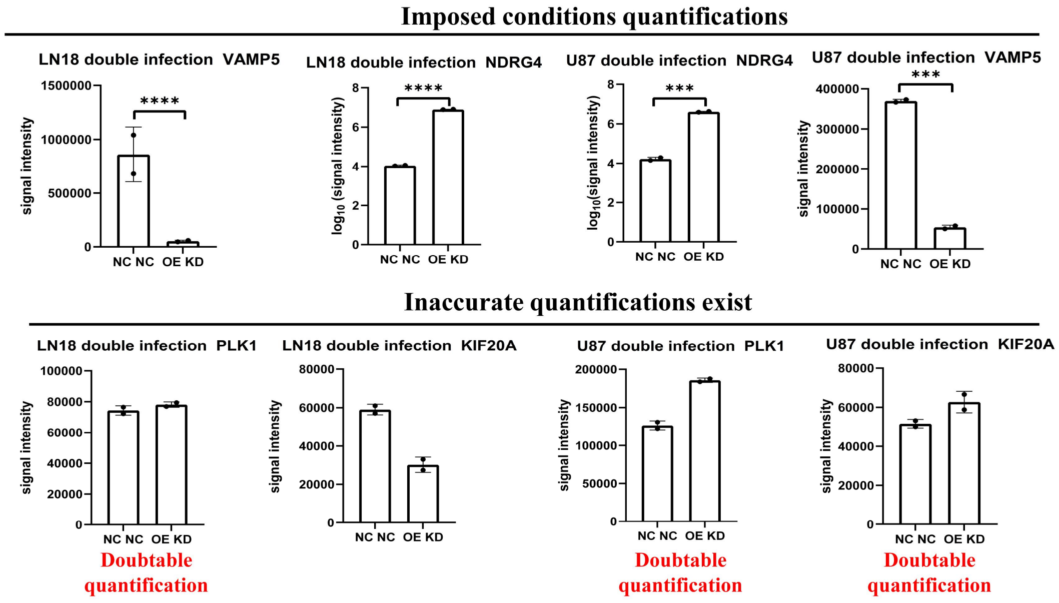


Fig S14: **Though the imposed conditions were correct, there existed inaccuracy of the quantification of proteomics results so more techniques should be applied to locate the possible diff-exp proteins.**

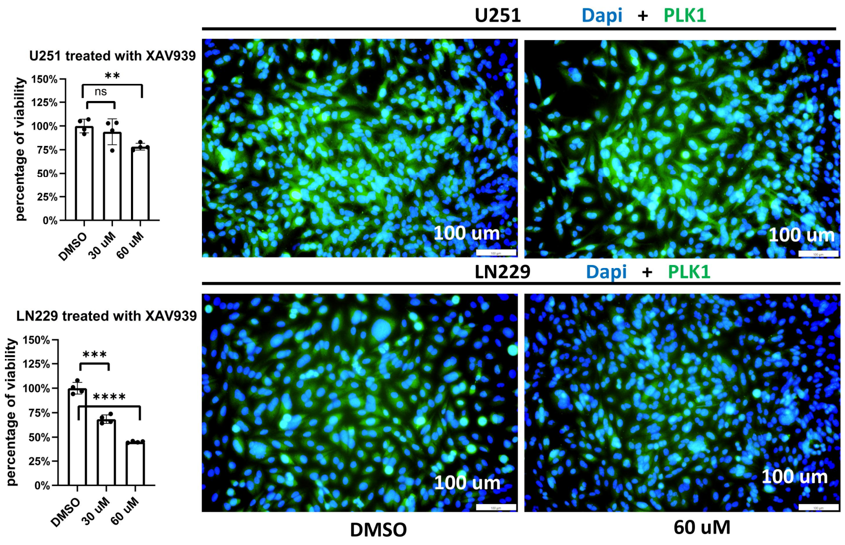


Fig S15: **Inhibiting TNKS independently may also suppress the cell growth without PLK1 down-regulations in some glioma cell lines indicating the inhibition mechanism parallel to PLK1.**

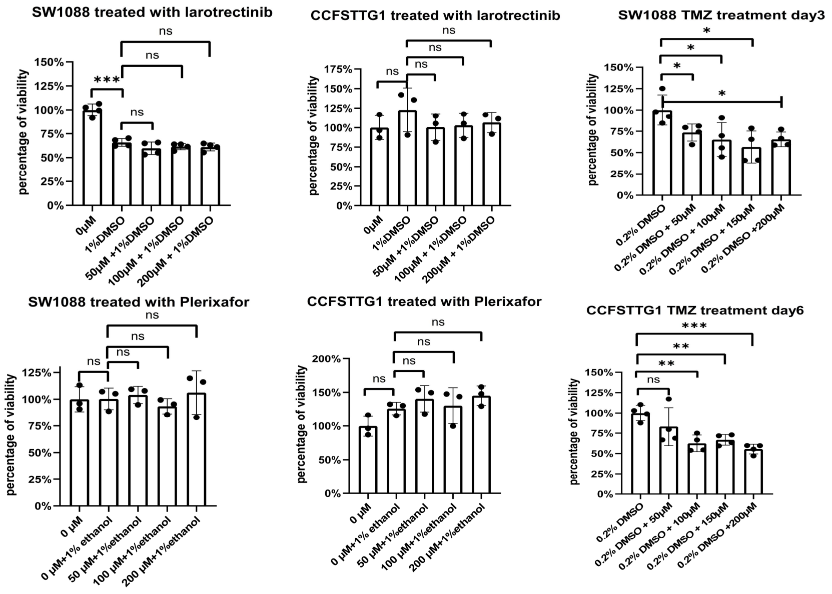


Fig. S16: **CXCR4 and NTRK3 (or their down-regulation) were validated not responsible for glioma growth (or growth inhibition) by using corresponding inhibitors, compared to positive control treated with TMZ.**
