## Supplementary material for "VAMP5 is a novel target for selective inhibition of gliomas with high NDRG4 expression via downregulating PLK1 and beyond": all supplemental tables

Table S1: Elements of SNARE gene set applied for GSEA in TCGA samples

| | | | | | | | | | | |
|---|---|---|---|---|---|---|---|---|---|---|
| **STX12** | **SNAP29** | **STX11** | **BNIP1** | **VAMP7** | **VAMP8** | **STX18** | **STX4** | **VTI1B** | **STX1B** | **STXBP3** |
| **STX12** | **STX6** | **STX16** | **STX3** | **SNAP47** | **YKT6** | **GOSR1** | **STX5** | **BET1** | **STX17** | **PRKCG** |
| **VAMP5** | **STX7** | **VAMP1** | **VAMP2** | **SEC22B** | **SNAP23** | **SNAP25** | **STX19** | **TSNARE1** | **NAPA** | **PRKCB** |
| **GOSR2** | **STX1A** | **VAMP3** | **STX10** | **VAMP4** | **BET1L** | **USE1** | **VTI1A** | **STX8** | **NAPB** | **PRKCA** |

Table S2: Sequences cloned into lentivirus vector. Red region is siRNA, and blue region is its anti-sense; Brown region is microRNA binding sequence.

| Lentivirus functions | Sequences 5—3 plus strand |
|---|---|
| Negative control LV | GATCCGTTCTCCGAACGTGTCACGTAATTCAAGAGATTACGTGACACG TTCGGAGAATTTTTTC |
| (TET-ON) VAMP5_KD LV | GATCCGATATGAGCTCAACCTTCAACAAGATTCAAGAGATCTTGTTGA AGGTTGAGCTCATATCTTTTTTG |
| VAMP5-6×His OE LV | ATGGCAGGAATAGAGTTGGAGCGGTGCCAGCAGCAGGCGAACGAGGT GACGGAAATTATGCGTAACAACTTCGGCAAGGTCCTGGAGCGTGGTGT GAAGCTGGCCGAACTGCAGCAGCGTTCAGACCAACTCCTGGATATGAG CTCAACCTTCAACAAGACTACACAGAACCTGGCCCAGAAGAAGTGCTG GGAGAACATCCGTTACCGGATCTGCGTGGGGCTGGTGGTGGTTGGTGT CCTGCTCATCATCCTGATTGTGCTGCTGGTCGTCTTTCTCCCTCAGAGC AGTGACAGCAGTAGTGCCCCACGGACCCAGGATGCAGGCATTGCCTCA GGGCCTGGGAACCATCATCACCATCACCAT |
| NDRG4-6×His OE LV | ATGCCGGAGTGCTGGGATGGAGGAGCCAAGAGCGGAGGCTG CCCAGAGTGTCCAGCACGGTCTCTCCCCTTCAGGAACATGACA TCGAGACACCCTACGGCCTTCTGCATGTAGTGATCCGGGGCTCC CCCAAGGGGAACCGCCCAGCCATCCTCACCTACCATGATGTGG GCCTCAACCACAAACTATGCTTCAACACCTTCTTCAACTTCGAG GACATGCAGGAGATCACCAAGCACTTTGTGGTGTGTCACGTGG ATGCCCCTGGACAACAGGTGGGGGCGTCGCAGTTTCCTCAGGG GTACCAGTTCCCCTCCATGGAGCAGCTGGCTGCCATGCTCCCCA GCGTGGTGCAGCATTTCGGGTTCAAGTATGTGATTGGCATCGG AGTGGGCGCCGGAGCCTATGTGCTGGCCAAGTTTGCACTCATC TTCCCCGACCTGGTGGAGGGGCTGGTGCTGGTGAACATCGACC CCAATGGCAAAGGCTGGATAGACTGGGCTGCCACCAAGCTCTC CGGCCTAACTAGCACTTTACCCGACACGGTGCTCTCCCACCTCT TCAGCCAGGAGGAGCTGGTGAACAACACAGAGTTGGTGCAGA GCTACCGGCAGCAGATTGGGAACGTGGTGAACCAGGCCAACCT GCAGCTCTTCTGGAACATGTACAACAGCCGCAGAGACCTGGAC ATTAACCGGCCTGGAACGGTGCCCAATGCCAAGACGCTCCGCT GCCCCGTGATGCTGGTGGTTGGGGATAATGCACCCGCTGAGGA CGGGGTGGTGGAGTGCAACTCCAAACTGGACCCGACCACTACG |

| | |
|---|---|
| | ACCTTCCTGAAGATGGCAGACTCTGGAGGGCTGCCCCAGGTCA<br>CACAGCCAGGGAAGCTGACTGAAGCCTTCAAATACTTCCTGCA<br>AGGCATGGGCTACATGCCCTCAGCCAGCATGACCCGCCTGGCA<br>CGCTCCCGCACTGCATCCCTCACCAGTGCCAGCTCGGTGGATG<br>GCAGCCGCCCACAGGCCTGCACCCACTCAGAGAGCAGCGAGG<br>GGCTGGGCCAGGTCAACCACACCATGGAGGTGTCCTGTTGA |
| Mir-1301-3p sponge LV | GAAGTCACTCCCGCCAGCTGCAATATACGAAGTCACTCCCGCCAGCTG<br>CAAACATCGAAGTCACTCCCGCCAGCTGCAATCTTCAGAAGTCACTCC<br>CGCCAGCTGCAA |
| Mir-12135 sponge LV | TTTACATGTAACCTTTATATACTTTACATGTAACCTTTAACATCTTTACA<br>TGTAACCTTTATCTTCATTTACATGTAACCTTTA |

Table S3: QPCR primers sequence for microRNA sponge sequence test

| Mir-1301-3p sponge sequence | Mir-12135 sponge sequence | Human β-actin |
|---|---|---|
| F-primer:<br>CGACAATCACGCGTGAAGTCA | F-primers:<br>AAGCCCGGTGCCTGAATCTA | F-primers:<br>TAGTTGCGTTACACCCTTTCTTG |
| R-primers:<br>TCCAGCTCGAGTTGCAGCTG | R-primers:<br>GAGGTTGATTGTTCCAGACGC | R-primers:<br>TCACCTTCACCGTTCCAGTTT |
